# RNA splicing dynamics in CD8 T cells uncovers isoforms that impact T cell-mediated cancer immunotherapy

**DOI:** 10.1101/2025.09.07.674706

**Authors:** Shay Tzaban, Priyanga Appasamy, Elad Zisman, Shiri Klein, Reyut Lewis, Houlin Yu, Akanksha Khorgade, Marc A. Schwartz, Moshe Sade-Feldman, Thomas Eisenhaure, Oren Parnas, Aron Popovtzer, Cyrille Cohen, Eric Shifrut, Aziz M. Al’Khafaji, Rotem karni, Galit Eisenberg, Nir Hacohen, Michal Lotem

**Affiliations:** The Lautenberg Center for Immunology and Cancer Research, The Faculty of Medicine Hebrew University of Jerusalem, Jerusalem, Israel; Center for Melanoma and Cancer Immunotherapy, Hadassah Hebrew University Medical Center, Jerusalem, Israel; Hadassah Cancer Research Institute, Hadassah Hebrew University Medical Center, Jerusalem, Israel; Sharett Institute of Oncology, Hadassah Medical Center, Hebrew University of Jerusalem, Jerusalem, Israel; Broad Institute of MIT and Harvard, Cambridge, MA, USA; Krantz Family Center for Cancer Research, Department of Medicine, Massachusetts General Hospital, Boston, MA, USA; Department of Biochemistry and Molecular Biology, IMRIC, Hebrew University-Hadassah Medical School, Jerusalem, Israel; Department of Genetics and the Institute for RNA Innovation, Perelman Medical School, University of Pennsylvania, Philadelphia, PA, USA; Laboratory of Tumor Immunology and Immunotherapy, The Goodman Faculty of Life Sciences, Bar-Ilan University, Ramat Gan, Israel; Gray Faculty of Medical and Health Sciences, Department of Pathology, Tel Aviv University; George S. Wise Faculty of Life Sciences, School of Neurobiology, Biochemistry and Biophysics, Tel Aviv University; Dotan center for Advanced Therapies, Tel Aviv Sourasky Medical Center

**Author notes:** **Correspondence:** Shay Tzaban – Michal Lotem –.

## Abstract

Immune checkpoint blockade has transformed cancer therapy, yet many patients fail to respond, and few new targets have emerged beyond PD-1 and CTLA-4. Alternative splicing dramatically diversifies the T cell proteome, but the functional roles of most isoforms remain unknown. Here we constructed the first single-cell splicing atlas of human CD8⁺ T cells, capturing dynamic isoform programs across activation and subset differentiation. This revealed distinct splicing footprints that refine conventional transcriptomic states and highlight receptor families with isoform-level regulation. To functionally interrogate these candidates, we developed SpliceSeek, a CRISPR-based pooled screening platform that perturbs splice sites to redirect isoform usage. Using SpliceSeek, we uncovered isoform-specific immune checkpoints whose perturbation enhanced effector function and tumor control, including the LRRN3-203 isoform, which augmented cytokine secretion and antitumor immunity in mice models. Together, our results establish alternative splicing as a targetable layer of immune regulation and demonstrate the potential of isoform-focused screening to expand the landscape of cancer immunotherapy.

## Introduction

Immune-checkpoint blockade has transformed the therapeutic landscape of malignancies such as melanoma and renal-cell carcinoma, producing durable responses in up to half of treated patients [1–4]. Yet a substantial fraction of malignancies remains refractory, and beyond PD-1 and CTLA-4 and emerging targets such as TIM-3 and LAG-3, few new modulatory receptors have reached clinical maturity [5–7]. This stagnation underscores a critical need for innovative immunotherapeutic targets that lie outside conventional paradigms.

Alternative splicing provides a powerful mechanism to expand the proteomic repertoire beyond gene number [8–9]. By altering mRNA splice patterns, alternative splicing can dynamically reshape protein function, subcellular localization and stability [10–12]. Although genome databases report hundreds of thousands of isoforms, the vast majority remain functionally uncharacterized, particularly in immune cells [13–14]. Early work by Lynch and colleagues first hinted that alternative splicing occurs broadly across the immune system, generating multiple isoforms of receptors, adaptors and transcription factors to fine-tune effector functions [15–16]. More recent analyses demonstrate that alternative splicing modulates entire receptor families and signaling cascades [17–18]. Intriguingly, receptor splice variants can exert agonistic or inhibitory effects distinct from their full-length counterparts. For example, a PD-1 isoform retaining 28 nucleotides of intron 2 (PD-1^28) more potently suppresses T cell activation than canonical PD-1 [19]. Yet systematic exploration of such isoforms as immunotherapeutic targets has been limited.

Here, we set out to chart the landscape of alternative splicing in activated human CD8⁺ T cells and to identify individual isoforms with costimulatory or agonistic potential absent from full-length transcripts. We first generated a high-resolution splicing atlas of naïve and effector CD8⁺ T cells, then developed “SpliceSeek,” a pooled CRISPR-based screen targeting splice sites across 150 immune-receptor genes. Top candidates were validated through in vitro assays of cytotoxicity and further assessed in vivo using humanized mouse models. Collectively, our work establishes a generalizable framework: starting from isoform discovery to mechanistic characterization and ending with in vivo validation, which enables the development of isoform-selective immunotherapies.

## Results

### Single-cell RNA sequencing reveals distinct CD8⁺ T cell subsets

In a previous study [20], a correlation was found between splicing patterns of naïve T cells in patients’ pre-treatment blood samples and a favorable response to immune checkpoint inhibitor (ICI) treatment, suggesting a connection between RNA splicing in lymphocytes and immune proficiency. However, the interpretation of these findings was limited by the use of short-read sequencing, which lacks the resolution to comprehensively capture the diversity of alternative splicing.

To address this limitation, we used a bi-modal sequencing strategy to profile RNA splicing in CD8⁺ T cells. We isolated CD8⁺ T cells from healthy donors (n = 3) and collected samples at 0, 1, 4, 24, 48, and 72 hours of activation. By integrating PacBio long-read sequencing for precise isoform resolution [21] with Illumina short-read sequencing for gene expression quantification, we generated a high-resolution atlas of dynamic RNA expression and splicing in activated CD8 T cells, offering a detailed view of splicing patterns at single-cell resolution (Fig. 1a). After filtering, we retained 11,495 cells with both gene-expression and long-read isoform data, identifying over 15,000 unique transcripts across 8,764 genes. Of these, 4,932 genes expressed only a single detectable isoform, while 3,832 genes showed two or more isoforms. We observed all major classes of alternative splicing (Extended Data Fig. 1a), with exon skipping accounting for 31% of spliced isoforms, alternative 3′ and 5′ splice-site usage 26%, alternative untranslated-region events 35%, and intron retention 9%, closely matching reported human transcriptome patterns [22,23]. Notably, roughly 2% of isoforms had no Ensembl annotation, underscoring novel splicing diversity in activated CD8⁺ T cells.

**Figure 1:**
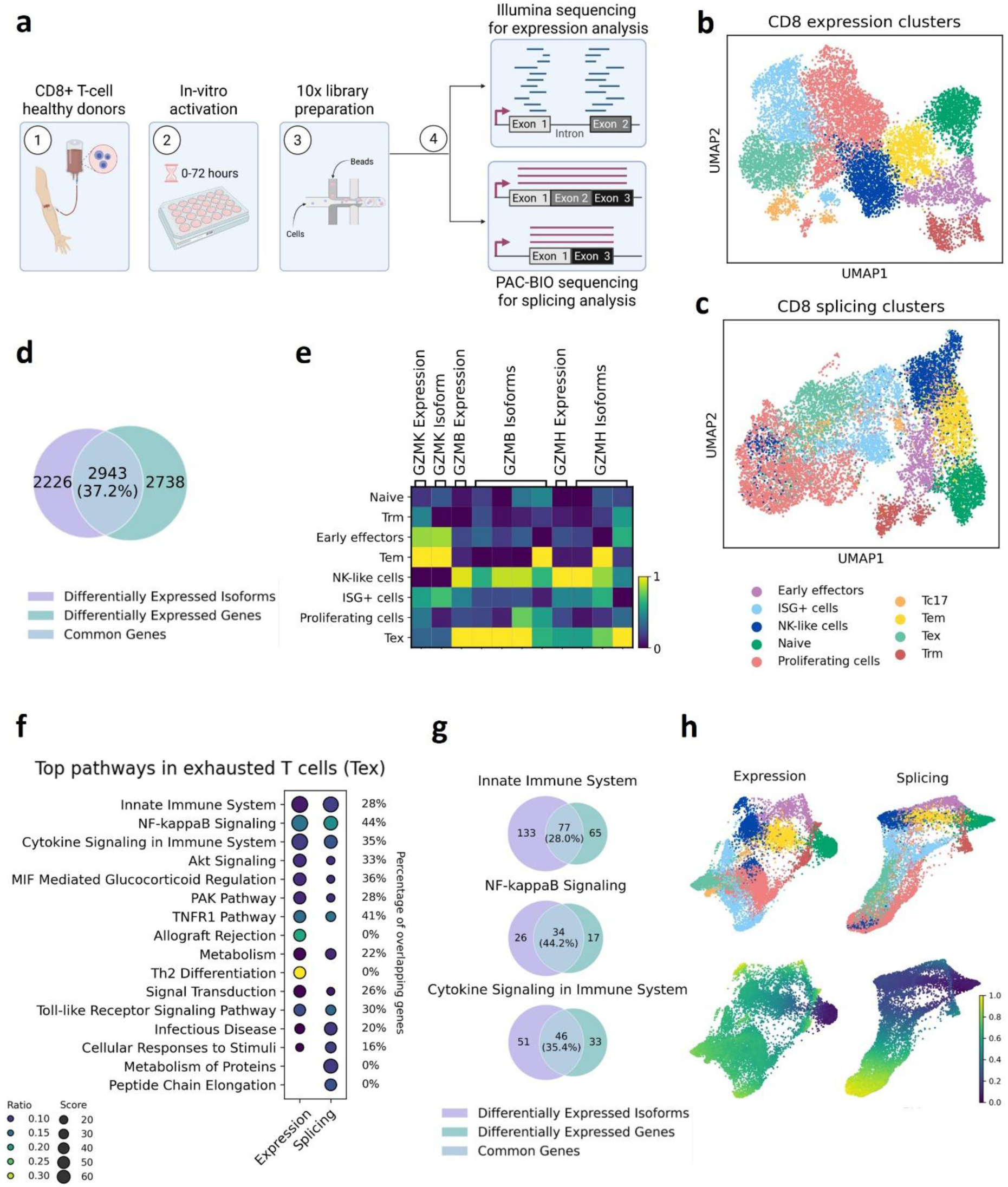
Single-cell atlas reveals gene expression–splicing interplay in CD8⁺ T cell activation and differentiation. **a,** Overview of the experimental design, Created in BioRender. **b-c,** UMAP projection of 11,495 CD8⁺ T cells collected from healthy donors (n = 3) at 0, 1, 4, 24, 48, and 72 hours of activation. Nine clusters were defined using gene- expression data (b) and then overlaid onto a UMAP constructed from splicing profiles (c). **d,** Venn diagram of differentially expressed splicing isoforms (purple) and genes (green) across all CD8⁺ T cell clusters; overlap (teal) denotes features that were significant in both analyses. **e,** Scaled heatmap of granzyme family gene-level and isoform-level expression across eight CD8⁺ T cell subsets. **f,** Gene set enrichment analysis of exhausted CD8⁺ T cells comparing splicing-based and expression-based datasets. Dot size represents the pathway enrichment score; dot color indicates the enrichment ratio; percentages at right side show the proportion of overlapping genes between splicing and expression for each pathway. **g,** Venn diagrams comparing differentially upregulated splicing isoforms (purple) and genes (green) in the top three enriched pathways. Overlap area shows shared genes. **h,** Force-directed graph of 11,495 CD8⁺ T cells derived from diffusion-map neighbor graphs. Cells are colored by clusters defined from gene-expression profiles (top left) and projected onto the long-read splicing-derived embedding (top right). Diffusion pseudotime trajectories are overlaid on force directed graphs (bottom row; scale 0–1, purple to yellow).

Unsupervised clustering of gene-expression data revealed 11 distinct CD8⁺ T cell clusters (Extended data 1b), which were consolidated into nine subsets based on gene signatures from previously published single-cell datasets [24–26] (Fig. 1b). A clear separation emerged between early (0, 1, and 4 hours) and late (24, 48, and 72 hours) activation time points (Extended data 1c). At early time points, five subsets were identified: naïve T cells, resident memory T cells (Trm), effector memory T cells (Tem), NK-like T cells, and early effector T cells. In contrast, later activation stages gave rise to four distinct subsets: exhausted T cells (Tex), TC17 cells, IFN-γ-stimulated T cells (ISG+), and proliferating CD8⁺ T cells. Notably, proliferating CD8⁺ T cells co-expressed naïve markers (CCR7) and ISG genes (STAT1) with high proliferative capacity (CENPV) but lacked cytotoxic markers (IFN-γ, GZMH), indicating an intermediate state between early effectors and exhausted cells (Extended Data Fig. 1d, table 1).

### RNA splicing defines CD8⁺ T cell subsets and reveals a continuum of differentiation states

We next asked whether T-cell clustering based on splicing patterns aligns with canonical subsets defined by gene expression. First, we performed unsupervised Leiden clustering of the splicing data (Extended Data Fig. 2a), which similarly distinguished early from late activation states (Extended Data Fig. 2b). We then quantified the concordance between splicing- and expression-derived clusters and found significant overlap (Adjusted Rand Index = 0.456, *p* < 0.001; Extended Data Fig. 2c). To assign biological identities to the splicing clusters, we mapped each Leiden splicing cluster to its corresponding expression-defined subset using cluster-overlap analysis (Extended Data Fig. 2d). Finally, we overlaid these assignments onto the splicing UMAP, demonstrating that splicing-based subpopulations closely recapitulate gene expression–defined T-cell states (Fig. 1c).

For example, the Tex population defined by expression corresponded to 84% of cells in splicing Leiden cluster 2 (Extended Data Fig. 2d). These cells were characterized at the gene-expression level by canonical exhaustion markers such as LAG3 and TIGIT, together with reduced expression of naïve-associated genes including TCF7 and LEF1. Importantly, our atlas now reveals splicing features unique to this state: isoforms of IL2RA, CCL3, and NKG7 were highly enriched in this cluster and provide the first splicing-level markers characteristic of Tex cells (Extended Data Fig. 2e).

Differential isoform expression analysis revealed that only ∼50% of differentially expressed isoforms (DEIs) originated from genes that were also differentially expressed at the gene level (DEGs) within the same cluster. The remaining ∼50% of DEIs reflected isoform-specific regulation without corresponding changes in overall gene expression. (Fig. 1d; Extended Data Fig. 2f, table 2). For example, although Granzyme H (GZMH) was differentially expressed in NK-like T cells, two of its isoforms were selectively enriched in other subsets (Tem and Tex). In contrast, both Granzyme K (GZMK) full gene transcript and its splice isoform showed concordant expression patterns, being upregulated in the same subsets (Fig. 1e). These findings demonstrate that isoform switching within granzyme genes can occur independently of changes in overall gene expression.

Recognizing that DEGs and DEIs only partially overlapped across clusters, we next performed pathway enrichment analysis separately for both DEGs and DEIs (Fig. 1f). While both DEGs and DEIs highlighted similar core immune pathways, including innate immunity, MHC class I antigen presentation, and cytokine secretion, the specific genes contributing to these pathways often differed. For example, in exhausted T cells, only 44% of NF-κB signaling genes were identified as both differentially expressed at the gene level and the isoform level; the remainder were unique to either DEGs or DEIs (Fig. 1g).

To dissect the respective contributions of gene expression and alternative splicing to CD8⁺ T cell differentiation, we performed diffusion-based pseudotime analysis to reconstruct cellular trajectories. Expression-based pseudotime primarily revealed a bifurcation between naïve and activated states. In contrast, splicing-derived pseudotime captured a continuous trajectory spanning from naïve cells through proliferative intermediates to exhausted (Tex) populations (Fig. 1h). Unlike RNA velocity approaches, which rely on spliced-to-unspliced transcript ratios to infer future states, our splicing-based pseudotime is derived solely from mature isoform profiles and does not incorporate unspliced read information. These findings highlight a coordinated yet distinct role for alternative splicing, which complements gene-level expression programs in guiding CD8⁺ T cell activation and fate decisions.

### Alternative splicing of immune-related genes characterizes the differentiation of exhausted **CD8⁺ T cells**

Given the contribution of RNA splicing to CD8⁺ T cell identity and function [27–29], we systematically examined splicing alterations in immune-related genes across differentiation states. This longitudinal approach enabled us to capture dynamic shifts in isoform usage associated with key functional transitions. Among these genes, PTPRC (encoding CD45), a well-characterized tyrosine phosphatase essential for T cell receptor (TCR) signaling, emerged as a model case. Previous studies [30–31] have highlighted the functional relevance of CD45 isoforms and their differential splicing across T cell subsets. While CD45 was broadly expressed at the gene level across all subsets, isoform-level analysis revealed eight distinct splice variants with subset-specific expression patterns (Fig. 2a). Notably, isoforms lacking exons 4–6 (CD45RO) were enriched in memory and activated subsets, consistent with their established prevalence in antigen-experienced T cells. Expression of these isoforms strongly correlated with HNRPLL, a splicing factor known to mediate CD45 exon exclusion [32] (Fig. 2b). This concordance with prior findings supports the robustness of our splicing-based profiling.

**Figure 2:**
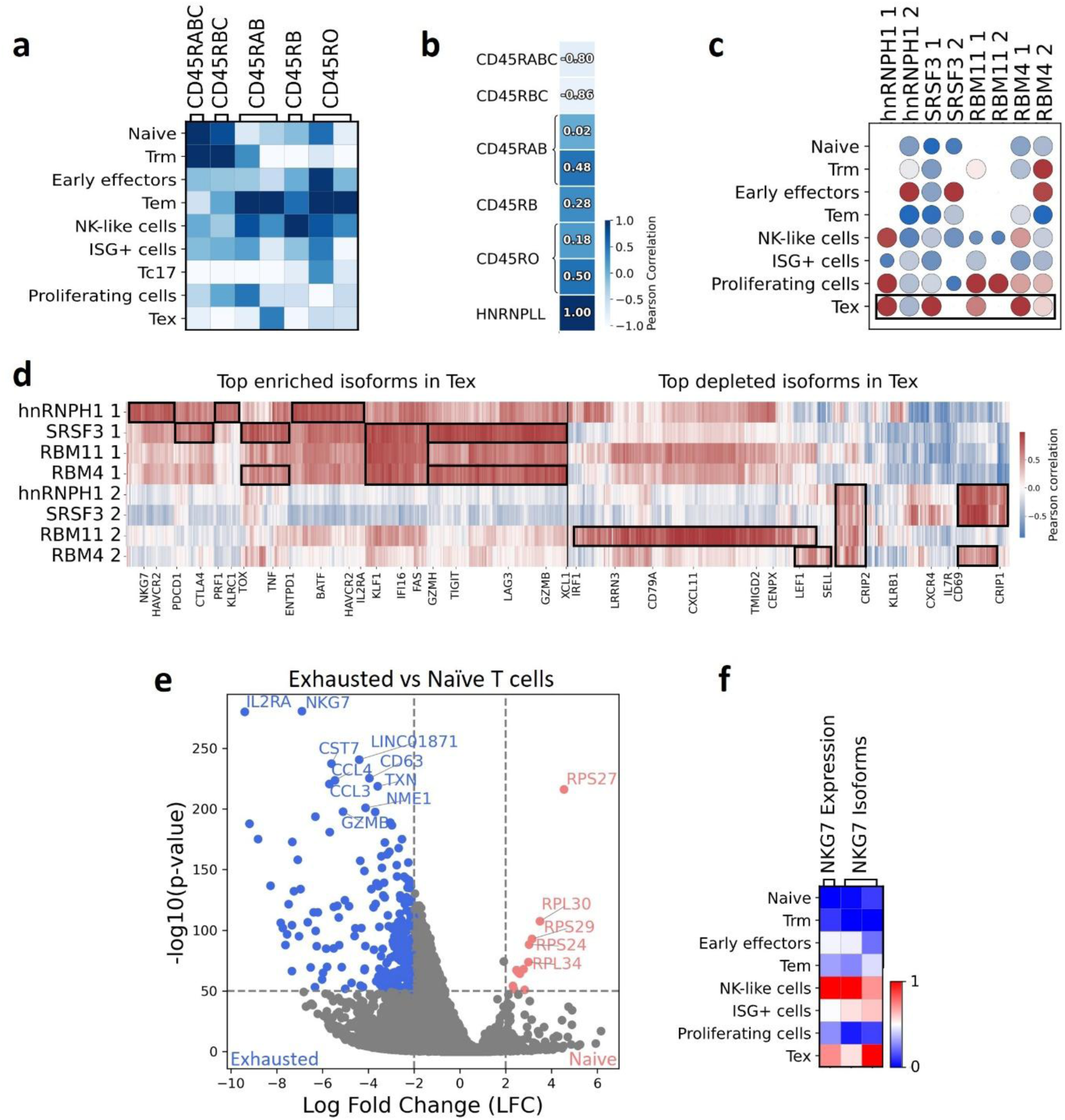
Alternative splicing of immune-related genes drives exhausted CD8⁺ T cell differentiation. **a,** Heatmap of expression of PTPRC (CD45) isoforms across CD8⁺ T cell subsets. Isoform abundances were min–max scaled per transcript (white = low expression; blue = high expression). **b**, Heatmap of Pearson correlation coefficients between expression of PTPRC (CD45) isoforms and the splicing regulator HNRNPLL across CD8⁺ T cell clusters. **c,** Dot plot of expression of splicing-factor isoforms across CD8⁺ T-cell subsets. Isoform abundances were min–max scaled per transcript (blue = low expression; red = high expression). **d,** Heatmap of Pearson correlation coefficients between splicing-factor isoforms (rows) and the top enriched (left) or depleted (right) isoforms in exhausted (Tex) CD8⁺ T cells (columns). Black boxes highlight clusters of isoforms showing coordinated correlation with each splicing-factor isoform. **e**, Volcano plot of differential isoform expression between exhausted (Tex) and naïve CD8⁺ T cells. Dashed lines mark thresholds; isoforms exceeding thresholds are blue (Tex) or red (naïve). **f,** Heatmap of NKG7 gene (left) and isoform (right) expression across CD8⁺ T-cell subsets; values are min–max scaled per transcript.

To uncover isoforms with potential functional relevance across CD8⁺ T cell states, we first looked for genes with multiple isoforms exhibiting distinct expression patterns across clusters. We reasoned that such patterns may reflect isoform-specific functions aligned with subset identity. This analysis revealed widespread isoform diversity, with notable enrichment in three functional categories: splicing regulators, immune receptors, and immune-related signaling molecules (Extended Data Fig. 3; Table 3). Several immune receptors showed isoform-specific enrichment in naïve, memory, or effector subsets, suggesting a role for alternative splicing in fine-tuning immune responses during differentiation.

Among these, isoforms enriched in the exhausted (Tex) subset were of particular interest due to their potential as immunotherapeutic targets. To investigate this further, we conducted a focused analysis of splicing factor usage and isoform regulation within Tex cells. Surprisingly, no splicing factor was significantly differentially expressed at the total gene level (Extended Data Fig. 4a). However, it is well established that splicing factors themselves are often regulated through alternative splicing [33–35], which can profoundly influence cellular phenotype. Indeed, several isoforms of key splicing regulators, including HNRNPH1, SRSF3, RBM11, and RBM4, were selectively enriched in Tex cells (Fig. 2c), while alternative isoforms of the same genes were enriched in distinct, non-Tex subsets.

To better understand the role of splicing factor isoforms in Tex cells, we quantified the Pearson correlation between the expression of each splicing factor isoform and that of Tex-DEIs, which are up- or downregulated in Tex cells compared to other subsets (Fig. 2d). We found that individual splicing factor isoforms enriched in Tex correlated with a distinct subset of upregulated DEIs (top left). In contrast, alternative isoforms of the same splicing factors, which were not enriched in Tex, correlated with different subsets of DEIs that were downregulated in Tex (bottom right). This divergent correlation pattern suggests functional specialization of splicing factor isoforms.

Building on the observed correlation between splicing factor isoforms and Tex-associated DEIs, we next sought to directly identify isoforms with functional relevance in the Tex population. Differential isoform analysis comparing Tex cells to other subsets revealed numerous Tex-enriched or depleted DEIs (Fig. 2e; Extended Data Fig. 4b-c). Among the most prominent were isoforms derived from IL2RA and NKG7, which differed between naïve and Tex subpopulations. While NKG7 is broadly recognized as a cytotoxic marker [36–38], we identified a previously uncharacterized, shorter isoform (NKG7-204) that was specifically enriched in Tex cells (Fig. 2f, Extended Data Fig. 4d–f). This Tex-specific isoform displayed an expression pattern that was distinct from the canonical gene and may possess unique functional properties. Together, these findings suggest that targeting alternatively spliced isoforms, rather than entire genes, may provide a new strategy for modulating T cell function.

### Establishing a CRISPR-Based Framework for Isoform Perturbation and Functional Screening in Primary Human T Cells

Our single-cell splicing atlas revealed extensive isoform diversity across CD8⁺ T cell states, highlighting alternative splicing as a potential regulatory layer in T cell identity. However, these insights remained correlative and lacked direct functional validation. To address this, we developed **SpliceSeek**, a CRISPR-based platform to selectively reprogram isoform usage in primary human T cells without altering coding sequences.

The platform integrates: (1) a CRISPR editing strategy that targets splice sites to rewire isoform expression (Extended Data Fig. 5a); (2) pooled screening in antigen-specific primary human T cells; and (3) a matched gene knockout arm to enable direct comparison between isoform-specific perturbation and full gene loss-of-function. Our longitudinal splicing analysis of CD8⁺ T cells revealed that immune receptors are among the most frequently affected by splicing alterations across different states and subsets. Based on this observation, we curated a focused library of 150 immunoglobulin superfamily (IgSF) receptors for functional screening, encompassing well-studied checkpoints such as PDCD1 (PD-1), HAVCR2 (TIM-3), and LAG3, alongside many with uncharacterized roles (Table 4).

To validate the feasibility of reprogramming splicing through CRISPR-mediated disruption of splice junctions, we used SLAMF6 as a test case. SLAMF6 is an immune receptor with three isoforms previously characterized in our lab [39]: the canonical form, a 3′ alternative splicing isoform, and an exon-skipping isoform. Targeting splice sites in intronic 5’ or 3’ junctions of Jurkat cells led to isoform switching without disrupting overall gene-level expression (Fig. 3a– c; Extended Data Fig. 5b-d), confirming the robustness of this approach at both the transcript and protein level. Notably, disruption of the 3′ splice site increased both exon skipping and 3′ alternative splicing, whereas disruption of the 5′ splice site selectively enhanced exon skipping without generating non-physiological isoforms.

**Figure 3:**
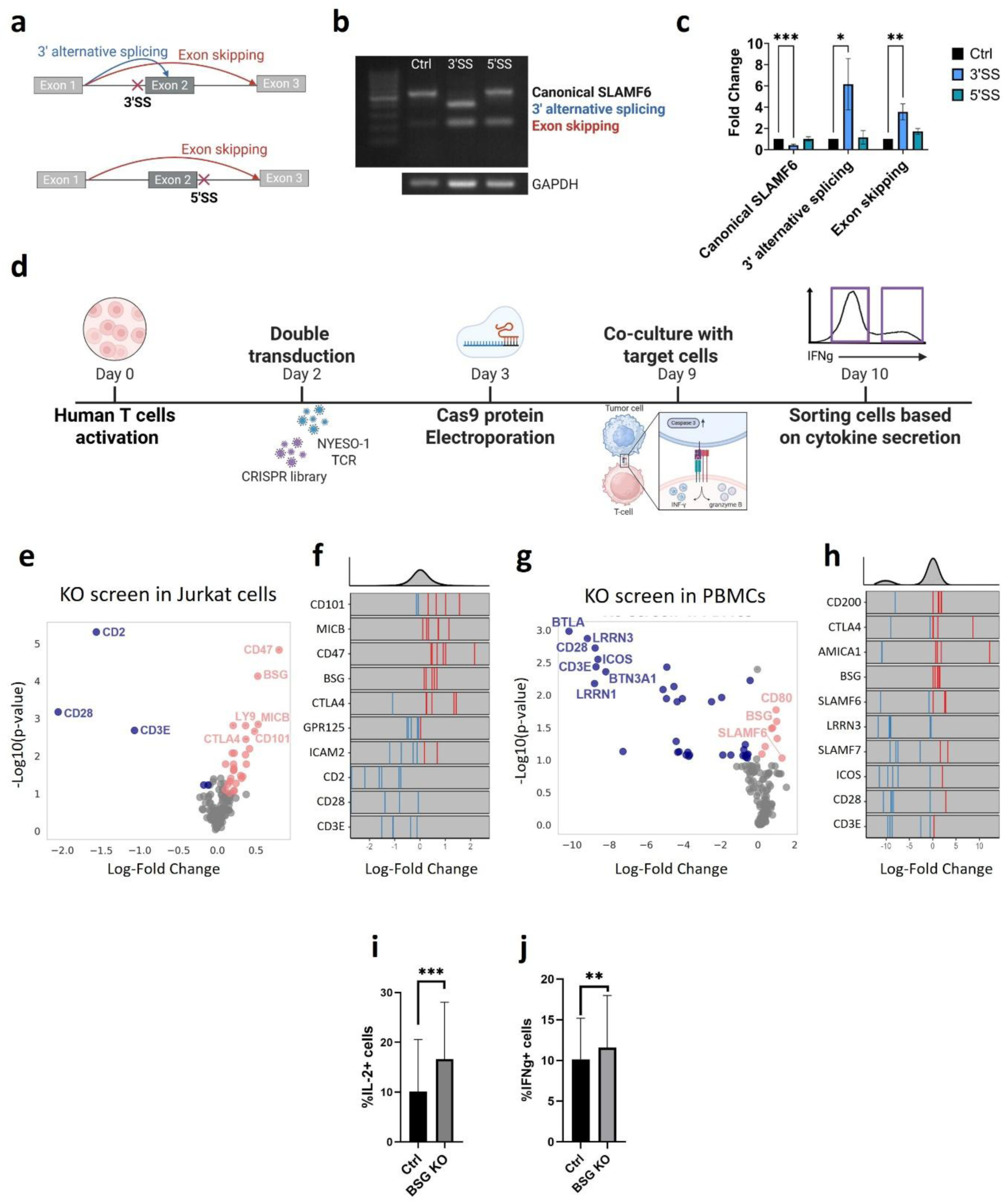
Establishing a CRISPR-Based Framework for Isoform Perturbation and Functional Screening in Primary T Cells. **a**, Schematic of CRISPR-Cas9 targeting of SLAMF6 splice sites. Disruption of the 3′ splice acceptor (3′SS; top) permits use of alternative downstream acceptor sites and exon skipping, whereas disruption of the 5′ splice donor (5′SS; bottom) yields exon skipping only. Created in BioRender**. b–c**, SLAMF6 isoform expression following splice-site editing in Jurkat T cells transduced with control, 3′SS or 5′SS guides. **b**, RT-PCR of SLAMF6 isoform and house-keeping control (GAPDH). **c**, RT-qPCR quantification of each isoform (mean ± SD; Ctrl n = 5, 3′SS n = 5, 5′SS n = 2). **d**, Schematic of the pooled SpliceSeek screen in primary, antigen-specific, human T cells. Created in BioRender. **e–f,** Reference gene knockout screen in Jurkat T cells activated by co-culture with T2 peptide loaded cell (n = 3 independent experiments). **e,** Volcano plot of sgRNA enrichment; red - guides enriched in IL-2-high cells; blue - guides enriched in IL-2-low cells. **f,** Density strips of guide-level log₂ FC for selected targets (top panel shows overall guide distribution). **g-h,** Reference gene knockout screen in primary human T cells (n=3 donors). **g,** Volcano plot of sgRNA enrichment; red - guides enriched in IFN-γ-high cells; blue - guides enriched in IFN-γ-low cells. **h,** Density strips of guide-level log₂ FC for selected targets (top panel shows overall guide distribution) **i,** Intracellular IL-2 production in Jurkat cells following overnight activation with PMA and ionomycin, comparing BSG knockout and control guides. **j,** Intracellular IFN-γ production in primary PBMCs after overnight co-culture with T2 cells, comparing BSG knockout and control guides. Data in **b** is representative of three independent experiments. P values were determined by mixed-effects model with Dunnett’s correction (**c**), and two-way ANOVA (**i,j**). *P < 0.05 **P < 0.01, ***P < 0.001.

For functional screening, we adapted a previously established CRISPR-based gene editing protocol for T lymphocytes [40]. Primary human T cells were engineered to express an NY-ESO-1:157-165/A2-restricted antigen-cognate TCR (NY-ESO-1 TCR), enabling antigen-specific T cell activation. Cells were transduced with the single-guide (sgRNA) library and electroporated with Cas9 protein under optimized conditions (Extended Data Fig. 5e–h). Following co-culture with peptide-pulsed T2 target cells, IFN-γ production, measured by intracellular cytokine staining (ICS), was used as both the functional readout and the sorting marker, shown to correlate with T cell cytotoxicity (Fig. 3d; Extended Data Fig. 5i–l).

In parallel, we performed a reference, gene-level loss-of-function CRISPR knockout screen using the same IgSF gene set, aiming to establish a functional activation threshold that their splicing isoforms must surpass. The screen was conducted with three T cell populations: (1) wild-type Jurkat cell line activated with PMA/ionomycin (Extended Data Fig. 6a-b), (2) Jurkat cells expressing the NY-ESO-1 TCR activated with NY-ESO-1:157-165 peptide (Fig. 3e-f), and (3) primary peripheral blood mononuclear cells (PBMCs) stimulated with antibodies against CD3 and CD28 (Fig. 3g-h). Jurkat cells were sorted based on IL-2 production, and PBMCs based on IFN-γ production, reflecting their dominant cytokine output.

Model-based Analysis of Genome-wide CRISPR/Cas9 Knockout (MAGeCK) revealed strong concordance between the two Jurkat activated populations (R = 0.42, p < 0.0001; Extended Data Fig. 6c) and substantial overlap with primary T cells (R = 0.33, p < 0.0001; Extended Data Fig. 6d), underscoring the consistency and generalizability of IgSF gene effects across distinct lymphocyte contexts. As expected, knockout of well-known co-stimulatory receptors (CD3E, CD28, ICOS) impaired effector function, while loss of the inhibitory receptor SLAMF6 enhanced IFN-γ production, confirming the detection capacity of the screen. Of interest, the reference screen revealed a strong agonistic effect following knockout of CD147 (BSG), a receptor known for its role in SARS-CoV-2 infection but not commonly associated with T cell function [41]. Single-guide validation confirmed increased IL-2 and IFN-γ production (Fig. 3k-l; Extended Data Fig. 6e-j). These results supported the sensitivity of our screening platform and offered functional context for interpreting isoform-specific effects in the SpliceSeek screen.

### SpliceSeek - High-throughput isoform perturbation reveals alternative splicing isoforms that enhance T cell effector function

Having validated our platform and established a functional reference for gene loss-of-function, we proceeded to implement the SpliceSeek screen. A pooled sgRNA library was constructed to target splice sites of the IgSF genes, guided by conserved splice junctions identified in our splicing atlas and public transcript annotations (RefSeq, Ensembl) [42–43]. The screen was performed on T cells from three healthy donors, engineered to express NY-ESO-1 TCR and stimulated with peptide-pulsed T2 cells. T cells were sorted into high and low IFN-γ–producing populations (Extended Data Fig. 7a), and 24 splicing perturbations emerged as enhancing IFN-γ production, among them splicing perturbation of LRRN3, SEMA4F and IL1R1 (Fig. 4a, Table 5).

**Figure 4:**
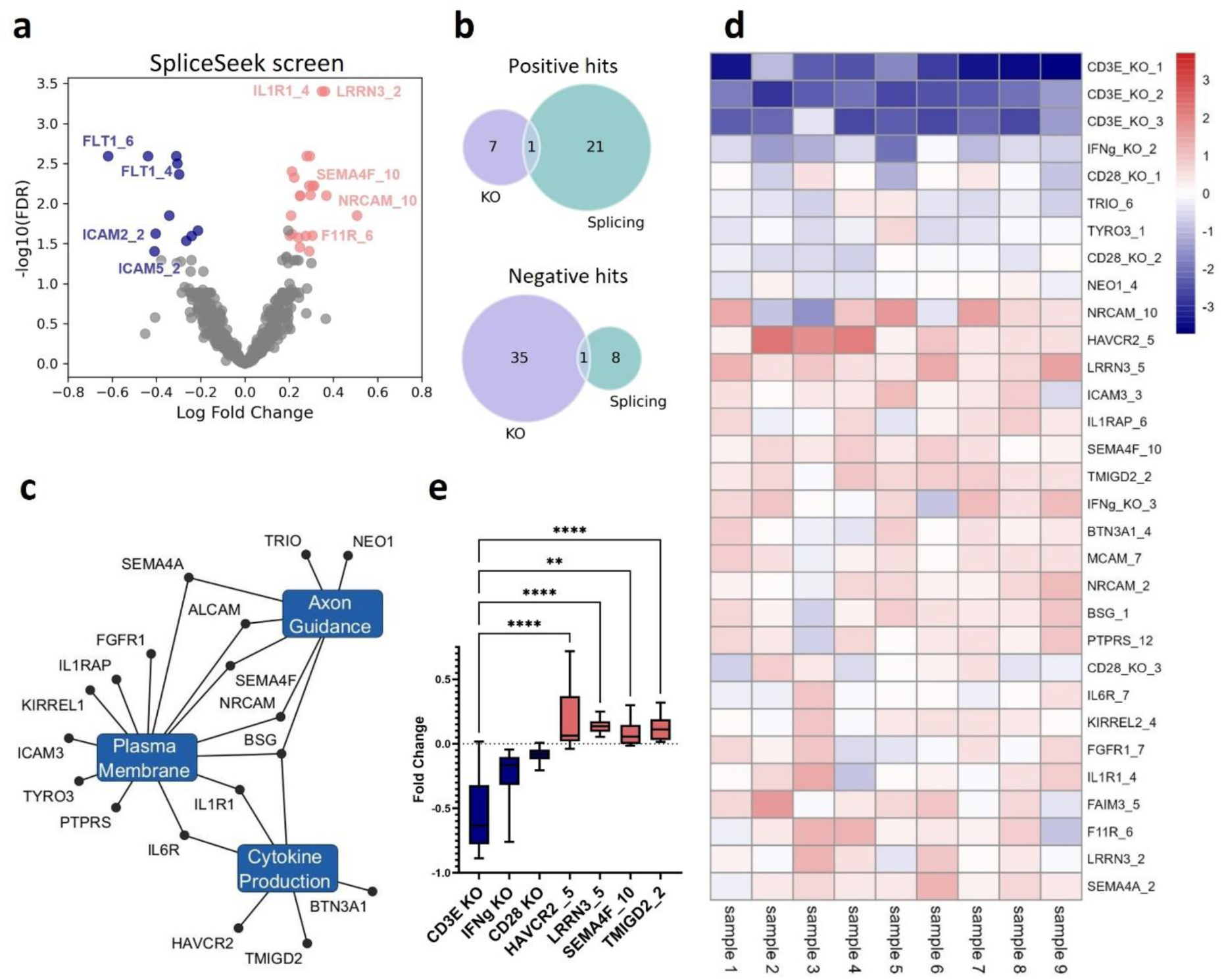
“SpliceSeek” - High-throughput isoform perturbation reveals alternative splicing isoforms that enhance T cell effector function: **a,** Volcano plot of the SpliceSeek CRISPR screen in primary T cells (n = 3 donors), showing sgRNA enrichment in IFN-γ– high (red) versus –low (blue) fractions. **b,** Venn diagrams comparing positive (top) and negative (bottom) hits between the SpliceSeek screen and the reference gene-knockout screen. **c**, Gene–Pathway network of SpliceSeek hits. Large Blue nodes represent pathways significantly enriched among the 24 positive hits, while smaller black nodes are individual isoform hits. **d,** Heatmap of log₂ fold-change for each sgRNA in the sub-library SpliceSeek Screen. Red indicates enrichment in the IFN-γ–high fraction; blue indicates depletion. **e**, Boxplots of sgRNA-level log₂ fold-change for the four top hits of sub-library versus negative controls. Sub-library experiments (d-e) were done in 3 human donors in three independent experiments. P values were determined by Friedman test with Dunn’s correction for multiple comparisons (e). **P < 0.01; ****P < 0.0001.

Comparison with the reference knockout screen revealed that most positive and negative hits identified by SpliceSeek were unique to splicing perturbation (Fig. 4b). Of the 24 positive hits, only BSG overlapped with the knockout screen, ranking among the top agonist receptors in both platforms. Unlike BSG, other isoforms like SEMA4F and NRCAM exhibited modest, non-significant effects in the knockout screen, yet demonstrated a markedly stronger impact when specifically perturbed at the isoform level. Moreover, LRRN3 and BTN3A1 exhibited distinct phenotype between the splicing and knockout screens; CRISPR knockouts of these genes reduced IFN-γ secretion, whereas targeted splicing perturbations of the alternative isoforms enhanced cytokine production (Extended Data Fig 7b).

Beyond their expected roles in membranal protein and regulation of cytokine production, pathway enrichment of these 24 positive hits revealed a striking overrepresentation of axon guidance and neuronal synapse pathways (Fig. 4c). In particular, members of the Semaphorin family (SEMA4F, SEMA4A), well-characterized synaptic adhesion molecules, surfaced as strong enhancers of IFN-γ output, despite little prior evidence for their function in T cells [44].

To validate the results of our screen, we performed a focused sub-library screen comprising the top 24 splicing isoforms hits from SpliceSeek, along with negative control genes such as CD3E. The screen was conducted in three independent experiments, each using T cells from three healthy donors, for a total of nine biological replicates. The top-performing splicing targets: HAVCR2, LRRN3, SEMA4F, and TMIGD2, consistently enhanced IFN-γ production across donors and experiments (Fig. 4d–e; Extended Data Fig 7c). To evaluate their impact on cytotoxicity, we conducted a parallel screen measuring CD107a mobilization and all four targets (hits) improved cytotoxic capacity (Extended Data Fig 7d). IFN-γ and CD107a readouts were strongly correlated (R = 0.63 Extended Data Fig 7e), supporting a coordinated enhancement of effector functions.

### Top splicing isoforms revealed by SpliceSeek reprogram transcript usage and enhance T cell effector functions

To validate the functional impact of individual isoform perturbations, we performed single-guide CRISPR editing of the top four gene isoforms identified as SpliceSeek hits - LRRN3, HAVCR2, SEMA4F, and TMIGD2 in primary human T cells. Following CRISPR editing, splicing shifts were confirmed by reverse transcription PCR (RT-PCR). Editing of LRRN3 led to a robust increase in the LRRN3-203 isoform (Fig. 5a–b, Extended Data Fig 8a–c), while SEMA4F editing elevated the expression of its corresponding isoforms, SEMA4F-201 (Fig. 5c-d, Extended Data Fig 8d–f). In contrast, the HAVCR2-207 isoform differed by only 37 base pairs from the canonical transcript, falling below our detection threshold (Fig. 5g, Extended Data Fig 8i). Editing of TMIGD2 resulted in a modest, non-significant increase in the TMIGD2-203 isoform. (Fig. 5e-f, Extended Data Fig 8g-h).

**Figure 5:**
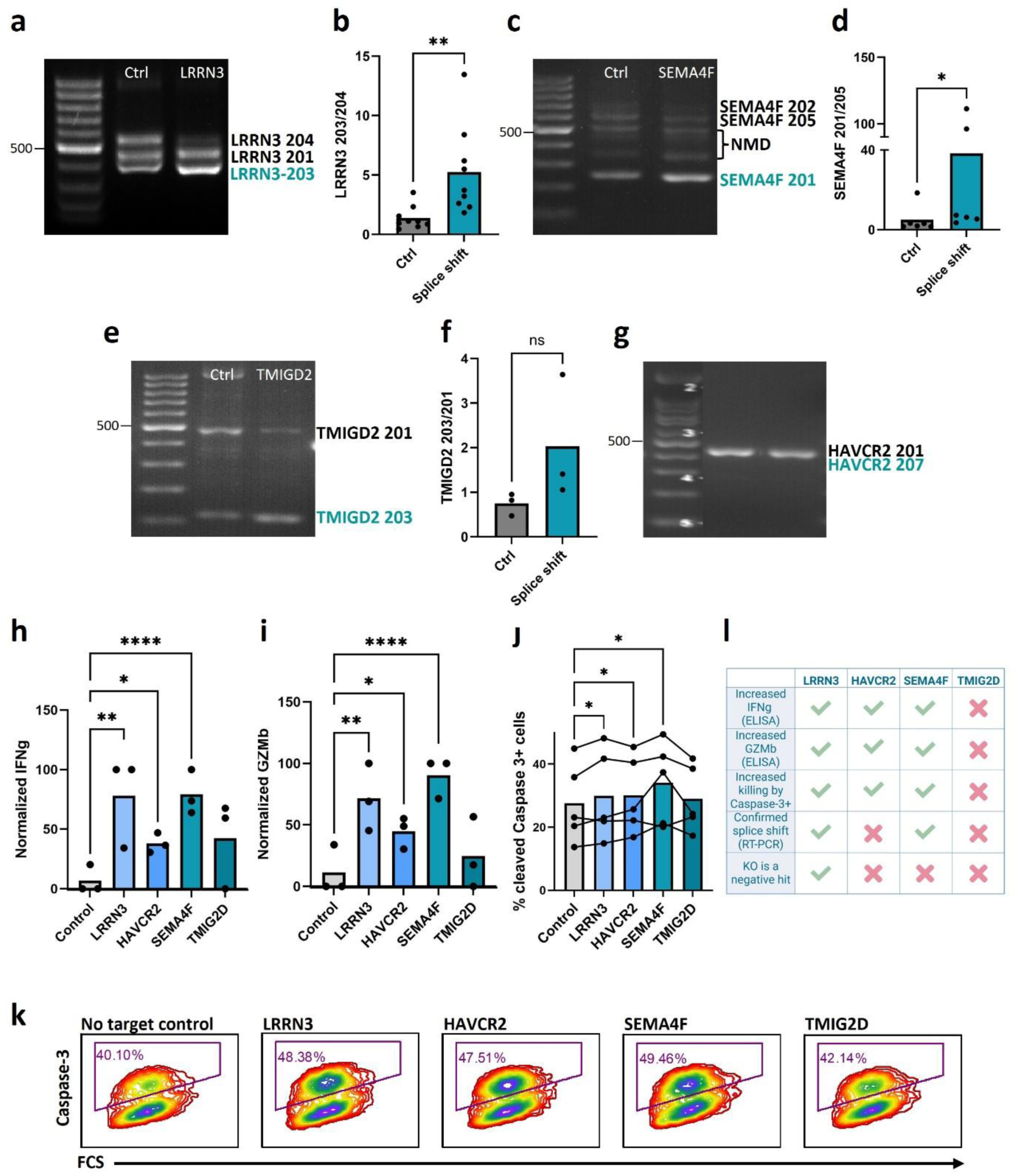
SpliceSeek hits reprogram transcript usage and enhances T cell effector functions. **a,** Representative RT–PCR of LRRN3 isoforms in cells transduced with non-targeting control (Ctrl) or LRRN3 splice-shift guide. **b**, Quantification of the LRRN3-203∶204 isoform ratio. **c,** Representative RT–PCR of SEMA4F isoforms in cells transduced with non-targeting control (Ctrl) or SEMA4F splice-shift guide. **d**, Quantification of the SEMA4F-205∶201 ratio. **e,** Representative RT–PCR of TMIGD2 isoforms in cells transduced with non-targeting control (Ctrl) or TMIGD2 splice-shift guide. **f**, Quantification of the TMIGD2-203∶201 ratio. **g,** Representative RT–PCR of HAVCR2 isoforms in cells transduced with non-targeting control (Ctrl) or HAVCR2 splice-shift guide**. h-i,** IFN-γ **(h)** and Granzyme B **(i)** secretion by ELISA in primary T cells transduced with control or splice-shift guides and stimulated overnight with 0.25 µg/mL anti-CD3/CD28. Values were normalized by min–max scaling of donor-wise mean measurements (bar show mean, dots as replicates, n = 3 donors, 3 different experiments). **j,** Cytotoxicity measured as the percentage of cleaved caspase-3–positive T2 target cells following 1.5 h co-culture with NY-ESO-1 TCR–expressing T cells edited with control or splice-shift. Bars show mean, dots represent five donors (each in 3 different experiment). **k,** Representative flow-cytometry contour plots of cleaved caspase-3 in T2 target cells after 1.5 h co-culture with T cells edited with non-targeting control or splice-shift guides. **l,** Summary of functional validation for the four top SpliceSeek candidates, Created in BioRender. All quantifications in panels b, d, and f were performed by densitometry in ImageJ on ≥3 biological replicates and are shown as mean with dots as replicates. P values were determined by paired Wilcoxon test (b, d, f); mixed-effects model with Dunnett’s multiple-comparisons test (I,j) and two-way ANOVA model with Dunnett’s multiple-comparisons test (h). *P < 0.05, **P < 0.01, ****P < 0.0001.

We next assessed the effector function of the manipulated isoforms in antigen-specific T cells. Splicing modulation of LRRN3, HAVCR2, and SEMA4F significantly enhanced the release of IFN-γ and granzyme B, whereas manipulation of TMIGD2 had minimal effect (Fig. 5h-i, Extended Data Fig 8j-k). Furthermore, NY-ESO-1 cognate lymphocytes engineered to express the selected isoforms induced increased caspase-3 cleavage in NY-ESO-1–pulsed T2 target cells, with the exception of TMIGD2, which showed no appreciable enhancement. (Fig. 5j-k).

Notably, in our reference knockout screen (Fig. 3g–h) LRRN3 knockout emerged as a top negative hit, significantly reducing IFN-γ production. In contrast, isoform-specific perturbation that promoted LRRN3-203 expression markedly enhanced cytokine secretion, revealing a reciprocal gain-of-function effect. Although isoform edits of HAVCR2 and SEMA4F also improved effector function, we prioritized LRRN3 for further validation due to its robust phenotype, reproducibility, and potential mechanistic relevance (Fig. 5l). Collectively, these results underscore the power of SpliceSeek to identify isoforms with intrinsic immunomodulatory activity and delineate LRRN3-203 as a potent enhancer of T cell reactivity.

### LRRN3-203 enhances protein level of LRRN3 gene

Leucine-rich repeat neuronal 3 (LRRN3) emerged from our SpliceSeek screen as a candidate modulatory checkpoint with an immune-regulatory role. LRRN3 is a type I transmembrane protein of the LRR family, comprising 11–12 extracellular leucine-rich repeats, an Ig-like C2 domain and a fibronectin type III domain [45]. It is best characterized in neural cells [46,47], but its role in T cells has not been defined. Recent studies identify LRRN3 as a marker of naïve and memory-like T cells [48–49] and link elevated expression to reduced T cell senescence [50]. Consistent with this, our single-cell atlas of human CD8⁺ T cells revealed preferential LRRN3 expression in naïve, proliferating, resident-memory and ISG⁺ subsets (Fig. 6a–b), suggesting a role in early activation states.

**Figure 6:**
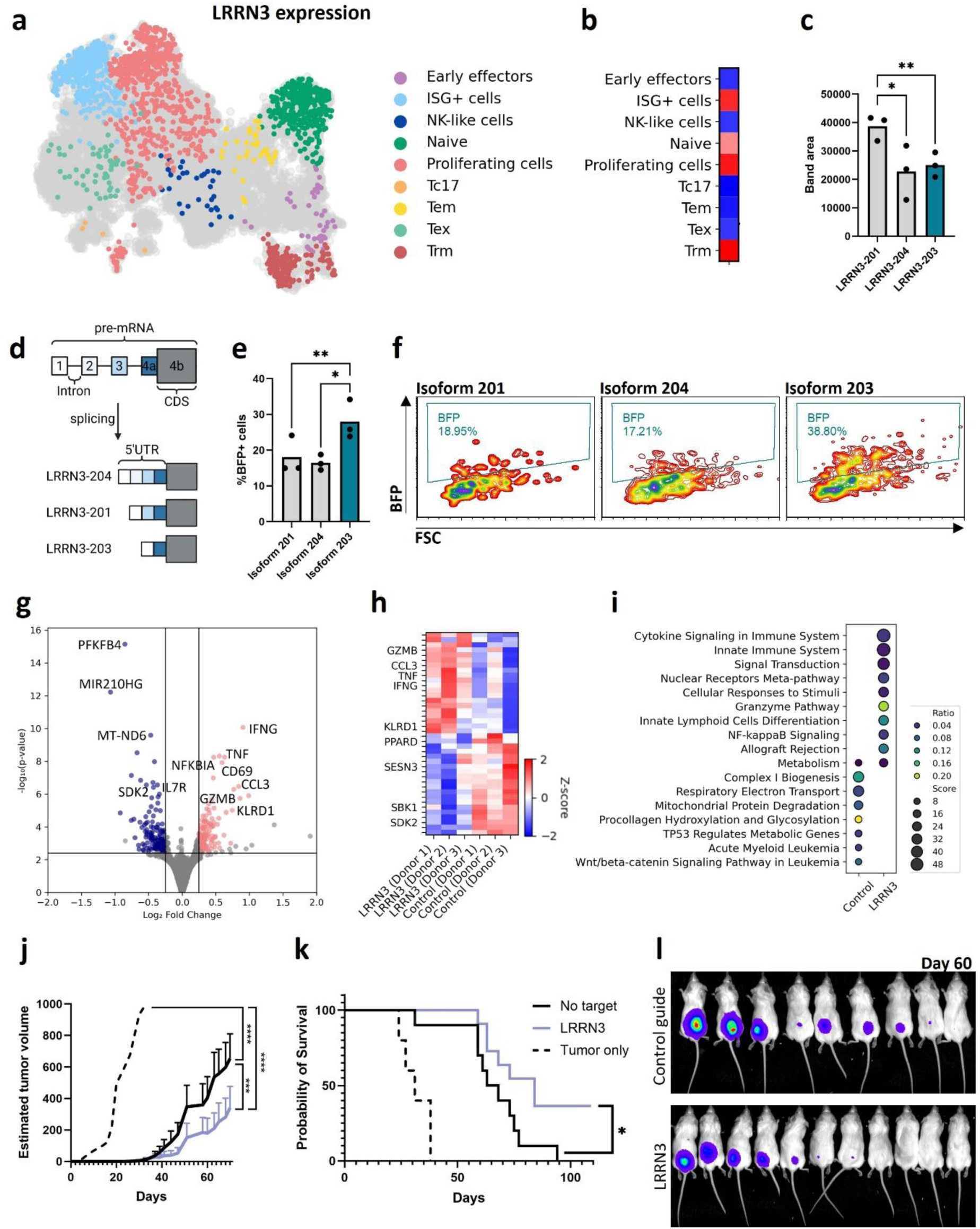
LRRN3-203 enhances protein expression and augments antitumor immunity. **a,** UMAP of CD8⁺ T cell expression from 11,495 cells showing LRRN3 transcript abundance by clusters. **b**, Heatmap of LRRN3 gene expression across CD8⁺ T cell subsets (blue = low expression; red = high expression). **c**, Densitometric quantification of RT-PCR band areas for LRRN3 isoforms in ImageJ. Band Area was measured for each isoform across three donors each in three independent experiments. Bars show mean, dots are replicates. **d**, Schematic of LRRN3 pre- mRNA and its three 5′-UTR isoforms, all encoding an identical protein. Created in BioRender. **e**, Jurkat cells transfected with BFP reporters bearing different LRRN3 5′-UTR isoforms (mean; n = 3 independent experiments). **f**, Representative flow-cytometry contour plots of Fig 6e. **g**, Volcano plot of DESeq2 differential expression between LRRN3-splice-shift and control T cells (n = 3 donors, red - up-regulated in LRRN3 cells, blue - down-regulated). **h**, Heatmap of Z-score–normalized expression for the top differentially expressed genes in LRRN3-shift versus control cells across donors. **i**, Gene-set enrichment analysis of pathways in LRRN3-shift versus control cells. Dot size represents the enrichment score, and dot color indicates the ratio of pathway genes detected. **j-l,** NSG mice were co-injected subcutaneously with luciferase-labeled A375 melanoma cells and human T cells engineered with either a non-targeting (No target), LRRN3 splice-modulating guides or no treatment control (No treatment n=5, No target n=10, LRRN3 n=11). **j**, Tumor volume over time (mean ± SEM). **k**, Kaplan–Meier overall survival curves. **l,** Representative bioluminescence images (day 60). P values were determined by two-way ANOVA with Tukey’s correction (d,j), one-way ANOVA with Dunnett’s correction (e), and log-rank test (k). *P < 0.05, **P < 0.01, ***p < 0.001, ****p < 0.0001.

RT-PCR of primary human T cells (Extended Data Figure 8a-c) detected three LRRN3 transcripts: LRRN3-201, -203 and -204, with isoform 201 being most abundant (Fig. 6c). These isoforms differ in their 5′ untranslated regions (UTR) yet encode an identical protein (Fig. 6d). Because CRISPR-mediated LRRN3 knockout reduced T cell activation in our knockout reference screen (Fig. 3g-h), we hypothesized that shifting splicing toward a shorter UTR might elevate LRRN3 protein levels and thereby enhance function. To test this, we cloned each UTR upstream of a BFP reporter and transduced Jurkat cells. The construct bearing the LRRN3-203 UTR drove the greatest expression level of total BFP - ∼28% versus 16% and 18% for the -204 and -201 UTRs, respectively (Fig. 6e-f).

### LRRN3-203 predicts favorable survival independently of CD3E

To assess the clinical relevance of LRRN3 splice variants, we queried The Cancer Genome Atlas (TCGA) pan-cancer RNA-seq cohort. Total LRRN3 and its individual isoforms showed distinct expression patterns across tumor types (Extended Data Figure 9a). We then focused on four well-annotated patient cohorts: lower-grade glioma (LGG), lung adenocarcinoma (LUAD), pancreatic adenocarcinoma (PAAC) and prostate adenocarcinoma (PRAD), and correlated expression with overall survival. While higher total LRRN3 was modestly associated with improved prognosis, this effect was amplified for the LRRN3-203 isoform and absent for LRRN3-201 (Extended Data Figure 9b–d). For example, PRAD patients with high LRRN3-203 levels exhibited >95% ten-year survival versus ∼60% in the low-expression group (log-rank p=0.046).

To determine whether LRRN3-203 abundance reflects more than just the presence of tumor-infiltrating lymphocytes, we calculated the Spearman’s correlation between LRRN3-203 and the pan-T cell marker CD3E, revealing only a modest association (R ≈ 0.40; Extended Data Figure 9e). Notably, CD3E expression alone did not stratify patient outcomes (Extended Data Figure 9f), indicating that LRRN3-203 carries prognostic information beyond lymphocyte abundance.

### LRRN3-203 reprograms effector functions and augments antitumor immunity

To investigate the mechanism underlying LRRN3-mediated functional enhancement, we performed bulk RNA-seq on PBMCs from three healthy donors transduced with either a non-targeting control guide or an LRRN3-203–promoting guide. Differential expression analysis identified IFN-γ, TNFa and GZMb among the top upregulated transcripts in the LRRN3-modulated cells (Fig. 6g–h, Extended Data Figure 10a). Gene set enrichment analysis revealed strong activation of NF-κB signalling, cytokine-mediated pathways and granzyme-driven cytotoxicity, concomitant with suppression of oxidative phosphorylation and other cellular stress response pathways (Fig. 6i, Extended Data Figure 10b–c).

Finally, to determine in vivo impact, we co-injected NSG mice with A375 melanoma cells and engineered T cells expressing NY-ESO-1 TCR with either LRRN3-203–promoting or control guides. Mice receiving LRRN3-203–biased T cells exhibited significantly delayed tumor growth (Fig. 6j, Extended Data Figure 10d) and a survival advantage: 25% of the LRRN3-203 cohort were long-term survivors versus 0% of the controls (log-rank p = 0.02; Fig. 6k). Concordantly, in vivo bioluminescence imaging of luciferase-expressing A375 tumors on day 60 showed that 55% of mice infused with LRRN3-203–biased T cells remained progression-free, compared with just 33% of control-guide recipients (Fig. 6l, Extended Data Figure 10e). Collectively, these findings establish LRRN3-203 as a tunable checkpoint whose splicing-dependent regulation can potentiate CD8⁺ T cell cytotoxicity.

## Discussion

In this study, we generated a high-resolution splicing atlas of primary human CD8⁺ T cells during activation by integrating long-read transcriptomics with single-cell RNA sequencing [21]. This atlas reveals a vast landscape of alternative splicing events affecting genes involved in immune receptor signaling, splicing regulation, and other immune-related functions. It captures isoform-level dynamics that complement conventional gene expression–based clustering and enhance pseudotime inference. Notably, splicing profiles more accurately reflected differentiation trajectories and provided additional resolution for both T cell subset delineation and pathway-level analyses.

Many differentially spliced isoforms mapped to clusters distinct from their canonical counterparts. For example, we identified NKG7-204, an uncharacterized variant selectively enriched in exhausted CD8⁺ T cells (Tex). NKG7 is a well-established cytotoxicity marker expressed by NK and NK-like T cells and correlates with improved tumor control [36–38], yet this specific isoform has not been previously described. Dissecting the functional contributions of NKG7-204 versus the canonical gene will require isoform-specific expression or targeted splicing modulation.

SpliceSeek addresses a critical gap in functional isoform discovery by enabling pooled CRISPR-Cas9 perturbation of intron-exon junction splice sites to shift isoform ratios in their endogenous genomic context. Unlike prior strategies, such as dCas13d-mediated scanning of splice-regulatory sites (Recinos et al. [51]), exon deletion screens that identify essential exons (Xiao et al. [52]), or isoform specific knockdown approaches that reduce transcript levels without reciprocally enhancing alternative variants (Schertzer et al. [53]), our platform simultaneously overexpresses selected alternative isoforms at the expense of the canonical transcripts and allows for functional screening. Moreover, by integrating SLICE-seq methods for pooled screen in primary T cells [40] with an NY-ESO-1 TCR knock in, we achieved scalable, antigen specific screening in primary human T cells, advancing beyond previous Cas9 and Cas13 modalities in immune cells [54–56].

Many genome-wide CRISPR screens target broad gene sets, which can inadvertently overlook immune receptors that are both mechanistically critical and clinically targetable. To address this, we deliberately focused our SpliceSeek library on a curated panel of 150 immune receptors, enabling deep interrogation of receptor specific splicing without diluting screening depth. Importantly, our platform is readily extensible to larger libraries; the primary constraint remains the manual design of splicing regulatory guides, which can be streamlined as predictive models of splicing regulatory elements improve.

Interestingly, some of the top hits were genes previously annotated as neuronal, with known or predicted roles in synaptic adhesion or signaling, but with little information on their role in T cells. Despite their annotation, these IgSF members were consistently expressed in both CD4⁺ and CD8⁺ T cells and were enriched for adhesive motives including leucine rich repeats, raising the possibility that they contribute to immune synapse formation or stabilization.

Our screen identified the LRRN3-203 isoform as a top candidate: CD8⁺ T cells enriched for this 5′UTR splice variant exhibited enhanced cytokine secretion and cytotoxicity in vitro and superior tumor control in vivo. Remarkably, CRISPR mediated knockout of LRRN3 abrogated T cell activation to a degree comparable to blockade of CD3/CD28 signaling, indicating that LRRN3 has the potential to function as a critical co-stimulatory receptor.

Mechanistically, LRRN3-203 arises from skipping of an alternative 5′UTR exon, which increases protein translation efficiency, a pattern mirrored in a prior large-scale analysis showing that ∼12% of human genes harbor alternative 5′UTR isoforms and that shorter UTR variants correlate with enhanced protein output [57]. LRRN3 is a marker of naïve and central memory T cells and is upregulated in smokers’ peripheral blood [58], yet its isoform specific roles were previously unexplored.

Finally, we observed that high LRRN3-203 expression correlates with improved overall survival in several cancer cohorts, albeit based on retrospective, correlative analyses. These findings raise the intriguing possibility that therapeutic modulation of LRRN3 splicing, such as shifting the isoform equilibrium toward LRRN3-203, could deliver dual benefits: enhancing tumor cell vulnerability and bolstering CD8⁺ T cell effector function. Future efforts to develop splice-modulating therapies that preferentially favor LRRN3-203 biogenesis may therefore provide a unified strategy to sensitize malignant cells while simultaneously empowering immune effectors, ultimately improving patient outcomes.

## Acknowledgments

This research was supported by Dr. Miriam and Sheldon G. Adelson Medical Research Foundation (AMRF), the Melanoma Research Alliance (MRA) and the European Union’s Horizon 2021 Research and Innovation program under grant agreement no. 101057250. The authors wish to acknowledge the devoted technical work of Inna Ben-David, Anna Kuznetz, Yael Gelfand, Shay Bamany and Dr Lola Weiss.

## Methods

### Mice

NSG (NOD-SCID IL2Rγnull) mice were purchased from The Jackson Laboratory. All experiments were performed with male 8- to 12-week-old mice. All experimental animal procedures were approved by the Institutional Animal Committee of the Hebrew University of Jerusalem.

### Isolation, Cryopreservation and culture conditions for Peripheral blood mononuclear cells from healthy donors

Peripheral blood mononuclear cells (PBMCs) were purified from buffy coats of healthy donors obtained from the Hadassah Blood Bank under Institutional Review Board approval. Cells (5 × 10^7) were frozen in human AB serum (bioIVT, HUMANABSRMP-HI-1) supplemented with 10% DMSO. After thawing, cells were cultured at a density of 2 × 10^6 cells/mL in LymphoONE T Cell Expansion Medium (Takara Bio, WK552S) supplemented with 5% human AB serum, 50 µM 2-mercaptoethanol, and 2.5% HEPES.

All procedures performed in this study involving human participants were in accordance with Hadassah Medical Center ethical approval 0661-23-HMO protocol. Informed consent was obtained from all human participants.

### Single-Cell RNA and Isoform Sequencing of Activated CD8⁺ T Cells

CD8⁺ T cells from three healthy donors were isolated using the CD8⁺ T Cell Isolation Kit (Miltenyi, 130-096-495). Cells were activated with Dynabeads Human T-Activator CD3/CD28 (Thermo Fisher, 11161D) at a 1:1 bead:cell ratio for 0, 1, 4, 24, 48, or 72 hours; beads were then removed magnetically. After activation, cells were blocked with Human TruStain FcX™ (BioLegend, 422301) for 10 minutes at 4°C, stained with the TotalSeq™-A Antibodies (BioLegend, 394601, 394603, 394605, 394607, 394611, 394623) for 30 minutes at 4°C, and washed three times in PBS containing 0.4% BSA. Cells viability and concentration were assessed by trypan-blue exclusion, and only samples with >85% viability proceeded to library preparation.

Approximately 3000 viable cells per sample were loaded onto the 10× Genomics Chromium to generate Single Cell 3′ libraries. Illumina libraries were sequenced on a Novaseq 6000 to a mean depth of ∼50,000 reads per cell. For full-length isoform sequencing, MAS-ISO libraries were prepared for PacBio as described by Aziz et al. [21]. Briefly, 3′ cDNA libraries were first amplified with Kapa HiFi Uracil+ ReadyMix, then purified with SPRIselect beads and Dynabeads kilobaseBINDER. After streptavidin purification, Bound cDNAs were released from beads by USER enzyme digestion. Each sample was divided into 15 tubes and subjected to a second PCR with the MAS-ISO primer pool. Reactions were then pooled and SPRI-cleaned, USER-digested, and circularized by HiFi Taq DNA Ligase. Final libraries were purified with AMPure PB beads, quantified by Qubit and Agilent Genomic DNA ScreenTape, and loaded onto the PacBio Sequel IIe for circular consensus sequencing.

### Processing of long-read single-cell RNA-seq data (MAS-ISO-seq)

Long-read output of PacBio sequencer generated via MAS-ISO-seq protocol as described above were processed using the Kinnex single-cell processing pipeline, following the standard workflow. Initial deconcatenation of the circular consensus sequences (CCS) was performed with skera split (v1.0.99) to segment the reads into individual cDNA molecules (s-reads). These segmented reads were then processed with lima (v4.0.0) to remove adapter sequences and orient reads in the 5’ to 3’ direction. Full-length reads containing both barcodes and poly(A) tails were extracted using isoseq-tag (v4.0.0). Next, isoseq-refine (v4.0.0) was used to identify and trim poly(A) tails, remove concatemers, and classify full-length non-concatemer (FLNC) reads. Finally, isoseq-correct (v4.0.0) was applied to correct cell barcodes and tag reads, enabling the generation of cell-resolved full-length isoform data for downstream analysis.

Barcode correction and PCR duplicate removal were further refined using isoseq3 (v4.0.0). Deduplicated reads were converted to FASTA format using samtools (v1.18) and aligned to the reference genome with minimap2 (v2.26) in splice-aware mode. IsoQuant (v3.3.1) was used for reference-guided isoform identification and quantification. The resulting output was parsed and converted into a single-cell isoform expression matrix for downstream analysis.

### Processing of Single-Cell RNA seq and analysis of short read and long read

Demultiplexing and alignment of short read Illumina data to the GRCh38 (hu38) reference genome were performed with Cell Ranger (10× Genomics) using default settings. Filtered feature-barcode matrices were imported into Scanpy [59], and cells with fewer than 200 detected genes, >15% mitochondrial reads, or genes expressed in fewer than three cells were excluded. The remaining count matrix was normalized to counts per ten thousand, log₁₊₁ transformed, and reduced by Principal component analysis (PCA). To mitigate donor-specific variation, Harmony batch correction was applied to the PCA embedding [60]. Uniform manifold approximation and projection (UMAP) was used for two-dimensional visualization. Leiden clustering was then executed at resolutions of 0.6 (short reads) and 0.5 (long reads).

Short-read clusters were annotated manually based on the top differentially expressed genes (identified with sc.tl.rank_genes_groups) and cross-referenced against published CD8⁺ T cell single-cell datasets [22–24]. Top markers per cluster were selected manually. Cells present in only one modality were discarded, yielding a final cohort of 11,495 cells profiled by both short- and long-read sequencing. Short-read cluster labels were overlaid onto the long-read UMAP to compare isoform- versus gene-level clustering. Cluster concordance between short- and long-read modalities was quantified using the Adjusted Rand Index (ARI). Differential expression analysis in long reads was conducted using Fisher’s exact test on isoform counts. Genes and isoforms exhibiting |log₂ fold change| > 1 and FDR < 0.05 were considered significant and overlaps between differentially expressed genes (DEGs) and isoforms (DEIs) were visualized for each cluster using Venn diagrams.

Diffusion pseudotime (DPT) analysis was performed in Scanpy. First, a force-directed graph was built using the Fruchterman–Reingold (‘fa’) layout. A root cell, selected from the naïve-like cluster, was defined as pseudotime zero. Pseudotime values for all cells were then computed with sc.tl.dpt and visualized by coloring the UMAP embedding according to each cell’s diffusion pseudotime. Gene set enrichment analysis (GSEA) was performed using the GeneAnalytics module (geneanalytics.genecards.org) of the GeneCards Suite [61]. Tex-specific gene lists (|log₂FC| > 1, FDR < 0.05) were submitted with default settings to query curated Super pathways and Gene Ontology.

To identify genes encoding multiple isoforms with cluster-specific expression, long-read differential isoform results (Fisher’s exact test, |log₂FC| > 1, FDR < 0.05) were first filtered to exclude the Tc17 cluster. For each gene represented by ≥2 significant isoforms, we tabulated the set of clusters in which each isoform was enriched. We then screened all pairwise (and, if needed, three-way) combinations of isoforms per gene, selecting those pairs whose cluster sets were not completely overlapping. That is, each isoform exhibited enrichment in at least one cluster not shared by its partner. Genes meeting this criterion were compiled into a final list of mutually exclusive isoform candidates (Supplementary Table 3).

### Cell lines sources and cultures

Cell line sources were as follows: Jurkat E6-1 (ATCC, TIB-152), HEK293 cells (CRL-1573), T2 (ATCC, CRL-1992), T2-RFP (generated with Incucyte® Nuclight NIR Lentivirus, Sartorius), A375 melanoma (ATCC, CRL-1619), A375-luc (a kind gift from Polina Stepensky). Jurkat, T2 and A375 cells were maintained in RPMI 1640 (Gibco) supplemented with 10% fetal bovine serum (FBS; Gibco), 100 U/mL penicillin, 100 µg/mL streptomycin, and 2 mM L-glutamine (Gibco). HEK293 cells were cultured in Dulbecco’s Modified Eagle Medium (DMEM; Gibco) containing 10% FBS, 100 U/mL penicillin, 100 µg/mL streptomycin, and 2 mM L-glutamine. All cell lines were incubated at 37 °C in a humidified atmosphere with 5% CO₂ and were routinely screened for mycoplasma contamination.

### Design of sgRNAs to Disrupt Splice Junctions

Single-guide RNAs (sgRNAs) were manually designed by extracting 50 bp of genomic sequence flanking each intron–exon boundary and scanning for upstream NGG PAM motifs. Guide sequences (20 nt) were chosen so that the predicted Cas9 cut site (3 bp upstream of the PAM) fell within 5 bp of the exon–intron junction, consistent with the 2–6 bp window essential for spliceosome recognition. Candidate guides were then evaluated with the Synthego Verify Tool, and only those with minimal predicted off-target activity were retained for downstream experiments.

### Plasmids cloning and Libraries preparation

sgRNAs for splice-junction disruption or gene knockout were designed as described above or selected using the Synthego Guide Design Tool and synthesized by HyLabs. For individual guides, each 20-nt spacer oligonucleotide pair was annealed, phosphorylated, and ligated into BsmBI-digested lentiGuide-Puro (Addgene #52963) using T4 DNA ligase; correct insertion was confirmed by Sanger sequencing. Pooled sgRNA libraries (knockout, splicing, and sublibraries) were synthesized as custom oligonucleotide pools by IDT, PCR-amplified with Gibson-compatible overhangs, and cloned into BsmBI-digested lentiGuide-Puro with NEBuilder^®^ HiFi DNA Assembly (NEB, E2621S). After ligation, libraries were transformed into electrocompetent bacteria by company protocol (Enduro cells, Lucigen, #60242-2). Plasmid DNA was extracted with NucleoBond Xtra Midi Kit (#MAN-740410.50). Libraries were sequence-verified by deep sequencing before lentiviral production.

### NY-ESO-1 TCR

The NY-ESO-1 TCR (1G4-α95:LY) engineered T cell was a kind gift from Cyrille Cohen’s lab [62].

### Retroviral production

PG13 packaging cells stably expressing the NY-ESO-1 TCR were seeded at a density of 5.5 × 10⁶ cells per T175 flask. After 48 h, the culture medium was replaced with fresh medium. At 96 h post-seeding, the cell supernatant was harvested and centrifuged at 400 × g for 10 min to remove cellular debris. The clarified supernatant was subsequently frozen and stored for later use.

### Lentiviral production

Lentiviral particles were generated by co-transfecting HEK293T cells with the transfer vector (Addgene #52962 or #52963), pMD2.G (Addgene #12259), and psPAX2 (Addgene #12260). Cells were seeded in 6-well plates at ∼60% confluence in 3 mL DMEM and transfected using Lipofectamine 2000 (Thermo Fisher, 11668019) in Opti-MEM: 1.25 µg transfer plasmid, 0.44 µg pMD2.G, 0.81 µg psPAX2, and 7.5 µL Lipofectamine 2000 per well (total volume 250 µL). Viral supernatants were collected 48 h post-transfection and used immediately or stored at – 80 °C.

### Jurkat Cell Transductions

Jurkat cells were first transduced with Cas9-expressing lentivirus (Addgene #52962) by spinfection: 0.5 × 10^6^ cells were resuspended in 500 µL RPMI 1640 containing 10 µg/mL polybrene and an equal volume of viral supernatant (1:1 ratio), then centrifuged at 900 × g for 60 min at room temperature. Following spinfection, cells were incubated at 37°C, 5% CO₂ and, 48 h later, selected with blasticidin (20 µg/mL; Sigma Aldrich, SBR00022) for five days.

Cas9-expressing cells were then transduced with lentivirus (Addgene #52963) encoding SLAMF6 splice-site guides (3′SS, 5′SS), BSG (CD147) Knockout guides or a non-targeting control using the same spinfection protocol. Forty-eight hours post-infection, transduced cells were selected with puromycin (10 µg/mL; Sigma Aldrich, P8833) for five days prior to downstream assays.

### RNA extraction, RT-PCR, and qRT-PCR

Total RNA was extracted using the Quick-RNA MiniPrep Kit (Zymo Research, R1054) and reverse-transcribed with the qScript cDNA Synthesis Kit (QuantaBio, 95047) according to the manufacturers’ instructions. For end-point reverse transcription PCR (RT-PCR), target amplicons were generated using Taq DNA Polymerase (PCR Biosystems, PB10.23-02), separated on agarose gels, and band intensities quantified in ImageJ. Quantitative real-time PCR was carried out on a ViiA 7 Real-Time PCR System (Thermo Fisher) with PowerUp SYBR Green Master Mix (Thermo Fisher, A25741). Reactions were run in triplicate, normalized to β-actin, and relative expression levels calculated by the ΔΔCₜ method. Primer sequences are listed in Table 6.

### Flow Cytometry Assays

Flow cytometric analyses were performed on a CytoFlex (Beckman Coulter) and cell sorting on an FACSAria III (BD Biosciences). All data were analyzed using FCS Express 5 Flow Research Edition (De Novo Software).

For cell-surface staining, 1×10^5^ CRISPR-edited Jurkat cells or CRISPR-edited PBMCs were washed once in PBS and incubated with primary antibodies: goat anti-SLAMF6 (R&D Systems, AF1908), PE anti-human SLAMF6 (BioLegend, 317207), PE anti-human TCR Vβ13.1 (Biolegend, 362409), APC anti-human TCR Vβ13.1 (Biolegend, 362407), FITC anti-human CD3 (Biolegend, 317305) or PE anti-human CD147 (BioLegend, 306211) each at a 1:100 dilution in PBS for 30 min at 4°C in the dark. After two PBS washes, cells stained with AF1908 were incubated with APC-conjugated anti-goat secondary antibody (Jackson ImmunoResearch, 705-136-147) at 1 µg/mL in PBS for 30 min on ice. Following a final wash, cells were resuspended in PBS and acquired and analyzed.

### Intracellular Cytokine Staining of BSG-Knockout Cells

BSG-edited primary T cells were co-cultured with Antigen-presenting T2 cells loaded with the NY-ESO-1_157-165_ peptide SLLMWITQV (0.1–0.5 µg/mL peptide, genscript) by incubation at 37 °C for 1h, at a 1:1 effector:target ratio for 6 h at 37 °C. Separately, BSG-edited Jurkat cells were stimulated with PMA (50 ng/mL, Sigma Aldrich, P1585) and ionomycin (1 µg/mL, Sigma Aldrich, I3909) for 12 h. In both assays, brefeldin A (BioLegend, 420601) was added after the first hour at a 1:1,000 dilution to block cytokine secretion. Following incubation, cells were washed and stained for surface markers (CD147, Vβ13.1) as described above, then fixed and permeabilized using the Perm/Fix kit (eBioscience, 88-8824-00). Intracellular staining was performed with APC anti–human IFN-γ (BioLegend, 502512) or P.B anti–human IL-2 (BioLegend, 500324) antibodies (1 µL per 100 µL Perm/Fix buffer) for 30 min at room temperature. After washing in Perm/Wash buffer, cells were resuspended in PBS and analyzed by flow cytometry on a CytoFlex instrument.

### Cleaved caspase-3 detection

T2-RFP cells were pulsed with 0.1–1 µg NY-ESO-1 peptide (as described above), and co-cultured with CRISPR edited TCR-transduced T cells (1:1 ratio) for 90 min. Cells were then fixed and permeabilized (BD Biosciences, 554714), stained with V450 Rabbit Anti-Active Caspase-3 (BD Biosciences, 560627) for 30 min at room temperature, washed, and analyzed.

### Granzyme B and CD107a Degranulation Assay

CRISPR-edited T cells (1×10^5^ per well) were co-cultured in triplicate with either A375 melanoma cells or NY-ESO-1–pulsed (0.1–1 µg/mL) T2 targets at a 1:1 effector:target ratio. Brefeldin A (1 µg/mL; BioLegend) was added at the start of the 6 h incubation at 37 °C. After 4 h, anti-human CD107a antibody (BioLegend, 328607) was added for the remaining 2 h to capture degranulation. At the end of the assay, cells were washed, fixed, and permeabilized using Perm/Fix kit. Intracellular staining for FITC anti-human Granzyme B (BioLegend, 372205) and APC anti-human IFN-γ (BioLegend, 502512) was performed following the manufacturer’s protocol.

### CRISPR Editing of Primary T Cells

Cryopreserved PBMCs were thawed and cultured for 48 h in T cell medium supplemented with 300 U/mL IL-2 and 1 µg/mL anti-CD3/anti-CD28. Activated cells (0.5 × 10^6^ cells/mL) were spin-transduced with lentiviral supernatant (encoding either individual CRISPR guides or pooled libraries) in the presence of 10 µg/mL polybrene by centrifugation at 900 × g for 1 h at room temperature. Twenty-four hours later, cells were electroporated with 5 µg recombinant Cas9 protein using the Amaxa human T cell Nucleofection kit (Lonza, VPA-1002, program T-007). Alternative electroporation programs and Cas9 concentrations were tested (Extended Data 5e–f). For co-transduction with the NY-ESO-1 TCR, retronectin-coated plates (Takara Bio) were first loaded with retroviral supernatant encoding the TCR, and PBMCs were then spin-transduced and Cas9-electroporated as described above. Twenty-four hours after electroporation, cells were subjected to puromycin selection (10 µg/mL) for 48 h.

### CRISPR splicing and knockout screens in T cells

CRISPR-edited Jurkat cells or primary PBMC-derived T cells harboring pooled sgRNA libraries were recovered after puromycin selection (10 µg/mL) for 2–3 days, then activated. PBMCs were stimulated either by co-culture with NY-ESO-1–pulsed T2 cells (0.1 µg/mL peptide) or with anti-CD3/CD28 antibodies (0.1 µg/mL each) for 6 h, whereas Jurkat cells were stimulated with PMA (50 ng/mL) and ionomycin (1 µg/mL) or by co-culture with NY-ESO-1–pulsed T2 cells (0.1 µg/mL peptide) for 12 h. One hour after stimulation began, brefeldin A was added at a 1:1,000 dilution to inhibit cytokine secretion. Following activation, cells were washed and surface-stained with FITC–anti–human CD3 (1:100 in PBS) for 30 min at 4 °C, washed again, then fixed, permeabilized (eBioscience Perm/Fix kit), and stained intracellularly with anti– human IFN-γ or anti–human IL-2 antibodies (1:100 in Perm/Wash buffer). Samples were maintained on ice and sorted for cytokine-positive populations on a BD FACSAria III.

After sorting, genomic DNA was extracted from fixed high- and low-cytokine-secreting cell populations using the Quick-DNA FFPE Miniprep Kit (Zymo Research; D3067) per the manufacturer’s protocol. DNA concentration was determined by Qubit dsDNA HS Assay (Thermo Fisher). For each sample, 1–2 µg of gDNA was split across multiple 100 µL Q5 High-Fidelity PCR reactions (NEB) supplemented with Illumina-compatible CRISPR sequencing adapters (Yau & Rana, 2018). PCR cycling was: 98 °C for 30 s; 20 cycles of 98 °C for 10 s, 60 °C for 20 s, 72 °C for 30 s; and a final 2 min extension at 72 °C. Reactions were pooled, purified using SPRI-select beads (Beckman Coulter; B23318), and libraries were quantified by Qubit and assessed on an Agilent Bioanalyzer. Final libraries were sequenced on an Illumina NextSeq 500.

Sequencing reads were demultiplexed and sgRNA count matrices generated using poolq3 with default settings. Differential guide and gene enrichment analyses were then performed with the MAGeCK [63] test module (default parameters), which applies median-ratio normalization to correct for library size differences. Importantly, treatment and control amples from each donor were analyzed as paired replicates to control for inter-donor variability. MAGeCK estimates variance by sharing information across sgRNAs and fits a negative-binomial model to assess whether individual guides are significantly enriched or depleted between treatment and control samples. sgRNAs are ranked by their NB-derived P-values, and gene-level selection is determined using the α-RRA algorithm to identify positively and negatively selected targets.

### PBMCs activation for ELISA assay

Primary CRISPR-edited T cells (top splicing hits or non-targeting controls) were recovered under puromycin selection for 2–3 days and then either (1) co-cultured with T2 antigen-presenting cells loaded with the NY-ESO-1_157–165 peptide (SLLMWITQV; GenScript) at 0.25–0.5 µg/mL at a 1:1 effector:target ratio or (2) stimulated with anti-CD3/CD28 antibodies (0.25 µg/mL each) for 24 h at 37 °C, 5% CO₂. Following incubation, culture supernatants were collected, and secreted IFN-γ and Granzyme B concentrations were measured by ELISA (R&D Systems, DY285B and DY2906) according to the manufacturer’s protocol.

### 5′ UTR–Driven BFP Reporter Assay for LRRN3 Translational Activity

Lentiviral reporter constructs encoding candidate LRRN3 5′ UTRs upstream of BFP (5′UTR– BFP–IRES–Puro) were synthesized by GenScript. Jurkat cells were spin-transduced with these vectors (0.5 × 10^6 cells in 500 µL RPMI + 10 µg/mL polybrene, 900 × g, 1 h, room temperature) and recovered overnight. Two days after infection, cells were selected with puromycin (10 µg/mL) for 2–3 days. BFP fluorescence was then measured by flow cytometry (CytoFlex, Beckman Coulter) to compare translational output driven by each 5′ UTR.

### Bulk RNA sequencing and analysis of LRRN3 manipulated T cells

PBMCs were edited with LRRN3 splice-shift guide or control guide as described above. On day 9 post thawing, cells were collected, and RNA was isolated using the Quick-RNA MiniPrep Kit (Zymo Research, R1054). The quality and integrity of total RNA was measured using TapeStation (Agilent Technologies) and total RNA sequencing libraries were constructed. Sequencing was performed on the Novaseq 6000 to a mean depth of 50 million reads per sample. Samples were aligned to the hg19 (b37) reference genome, and counts were quantified using featureCounts. Raw read counts were normalized, and differential gene expression was quantified with DESeq2, with design accounting for donor variations. Log fold change larger than 0.25 and a false discovery rate cutoff of 10% was used to select significantly up and down regulated genes. Gene set enrichment analysis was performed using the GeneAnalytics module of the GeneCards Suite.

### TCGA Alternative Splicing Analysis

Alternative splicing and isoform-level expression of LRRN3 across TCGA cohorts were analyzed using the GEPIA2 web server (http://gepia2.cancer-pku.cn)[64]. Transcript-level abundances in tumor versus matched normal samples were retrieved via the “Expression DIY” module. Co-expression between LRRN3 and CD3E across cancer types was assessed using the Correlation Analysis tool. Kaplan–Meier survival curves stratified by median LRRN3 isoform expression were generated with the “Survival” module. All plots and data tables were exported directly from the GEPIA2 interface.

### Winn assay

Human PBMCs from two healthy donors were double-transduced with either an LRRN3- targeting sgRNA or a non-targeting control and with the NY-ESO-1 TCR. Transduction efficiencies were matched across groups by flow cytometry. After recovery and antibiotic selection, edited T cells were washed and combined with luciferase-expressing A375 melanoma cells at a 1:0.5 effector:target ratio (1 × 10^6^ A375-luc cells with 0.5 × 10^6^ TCR+ T cells) in a final volume of 100 µL PBS, then immediately injected subcutaneously into the dorsal flank of 8–10-week-old NSG (NOD-SCID IL2Rγnull) mice (n = 10 for control, n = 11 for LRRN3-edited and n=5 for no treatment control). Tumor size was measured in two perpendicular diameters three times per week. Mice were sacrificed when tumors reached 15 mm diameter in one dimension or when the lesion necrotized. Tumor volume was calculated as L (length) × W (width)^2^ × 0.5. On day 60 post-injection, mice received an intraperitoneal dose of D-luciferin (100 mg/kg), and bioluminescent signals were captured using an IVIS Spectrum imaging system to assess tumor burden.

## Supplementary Information

**Table S1.** DEGs by clusters

**Table S2.** DEIs by clusters

**Table S3.** mutually expressed isoforms within clusters

**Table S4.** List of IgSF receptors in library

**Table S5.** SpliceSeek results

**Table S6.** Primers list

**Supplementary Figures 1-9.**

**Extended Data Fig. 1:**
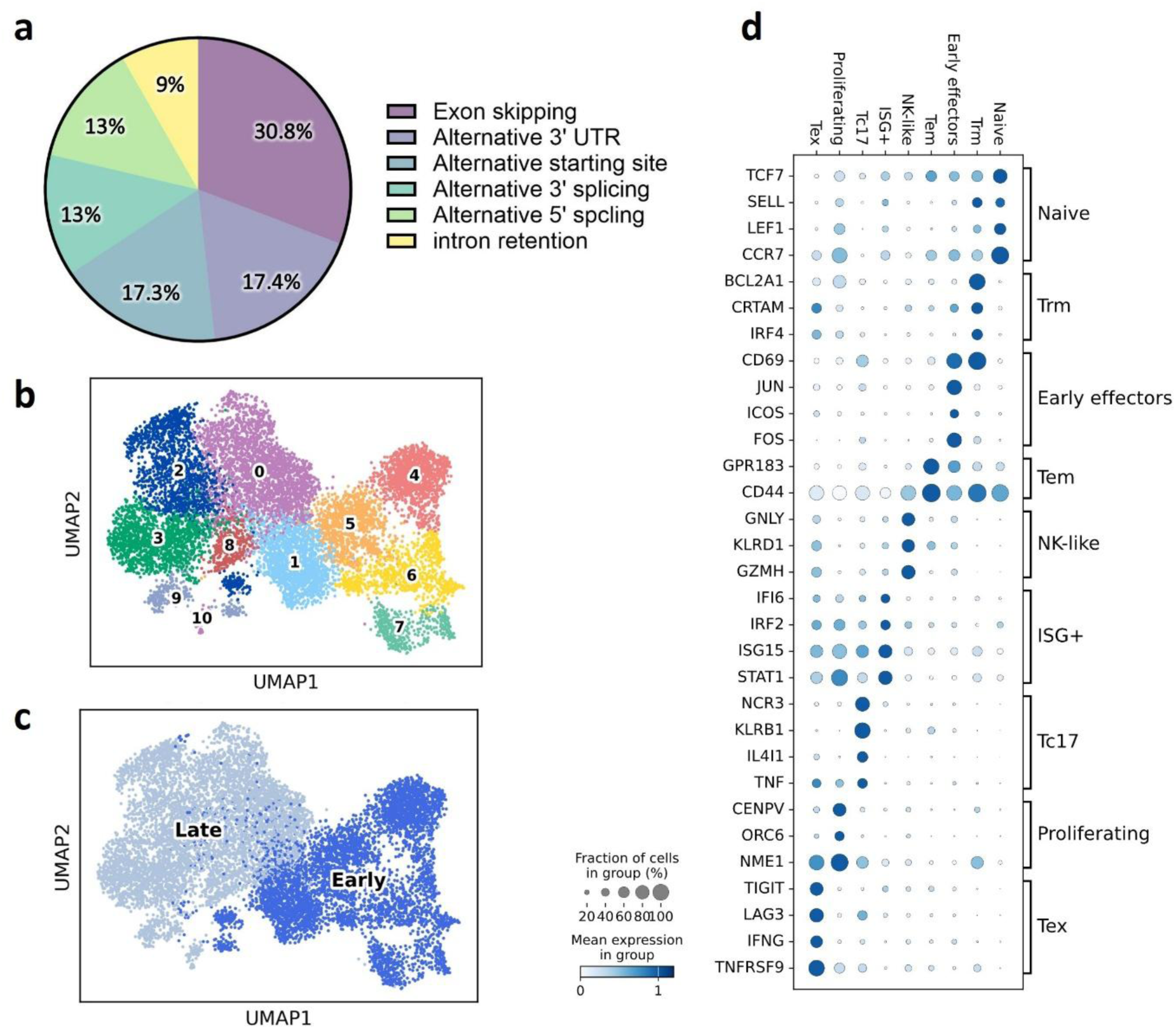
Clustering of CD8+ T cells based on expression data. **a,** Pie chart illustrating the relative prevalence of alternative splicing event types in human CD8⁺ T cells, based on long-read isoform analysis. **b,** UMAP embedding of 11,495 CD8⁺ T cells sampled from three healthy donors over six activation time points (0, 1, 4, 24, 48 and 72 h), with cells colored by unsupervised Leiden clusters based on expression data. **c,** UMAP of the same cells shown in b, colored by activation stage: early (0–4 h) versus late (24–72 h). **d,** Dot plot illustrating expression of cluster-defining genes across nine CD8⁺ T cell subsets. Dot size indicates the fraction of cells within each cluster expressing the gene, while color intensity reflects the mean expression level.

**Extended Data Fig. 2:**
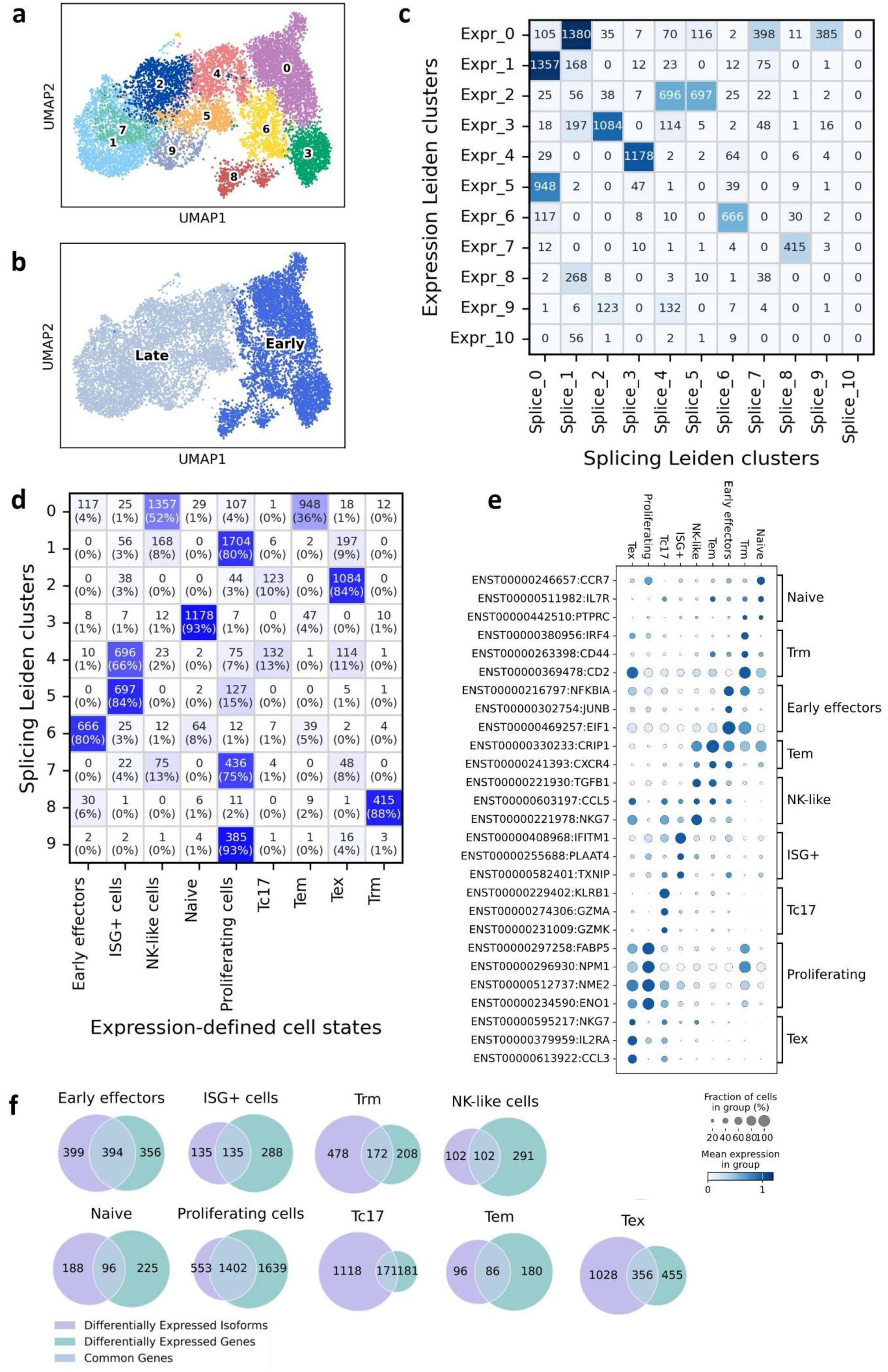
Clustering of CD8+ T cells based on splicing data. **a,** UMAP embedding of 11,495 CD8⁺ T cells sampled from three healthy donors over six activation time points (0, 1, 4, 24, 48 and 72 h), with cells colored by unsupervised Leiden clusters of splicing data. **b,** UMAP of the same cells shown in a, colored by activation stage: early (0–4 h) versus late (24–72 h). **c**, Confusion matrix comparing Leiden unsupervised cluster assignments from short-read expression (rows, Extended Data fig 1b) versus long-read splicing (columns, Extended Data fig 2a). Each cell shows the number of cells shared between the corresponding pair of clusters. **d**, Cluster-overlap analysis linking splicing- and expression-derived subsets. Heatmap shows the distribution of expression-defined T-cell states (columns) within each Leiden splicing cluster (rows). **e,** Dot plot illustrating expression of cluster-defining isoforms across nine CD8⁺ T cell subsets. Dot size indicates the fraction of cells within each cluster expressing the gene, while color intensity reflects the mean expression level. **f,** Venn diagram of differentially expressed splicing isoforms (purple) and genes (green) across all CD8⁺ T cell clusters; overlap (teal) denotes features significant in both analyses.

**Extended Data Figure 3:**
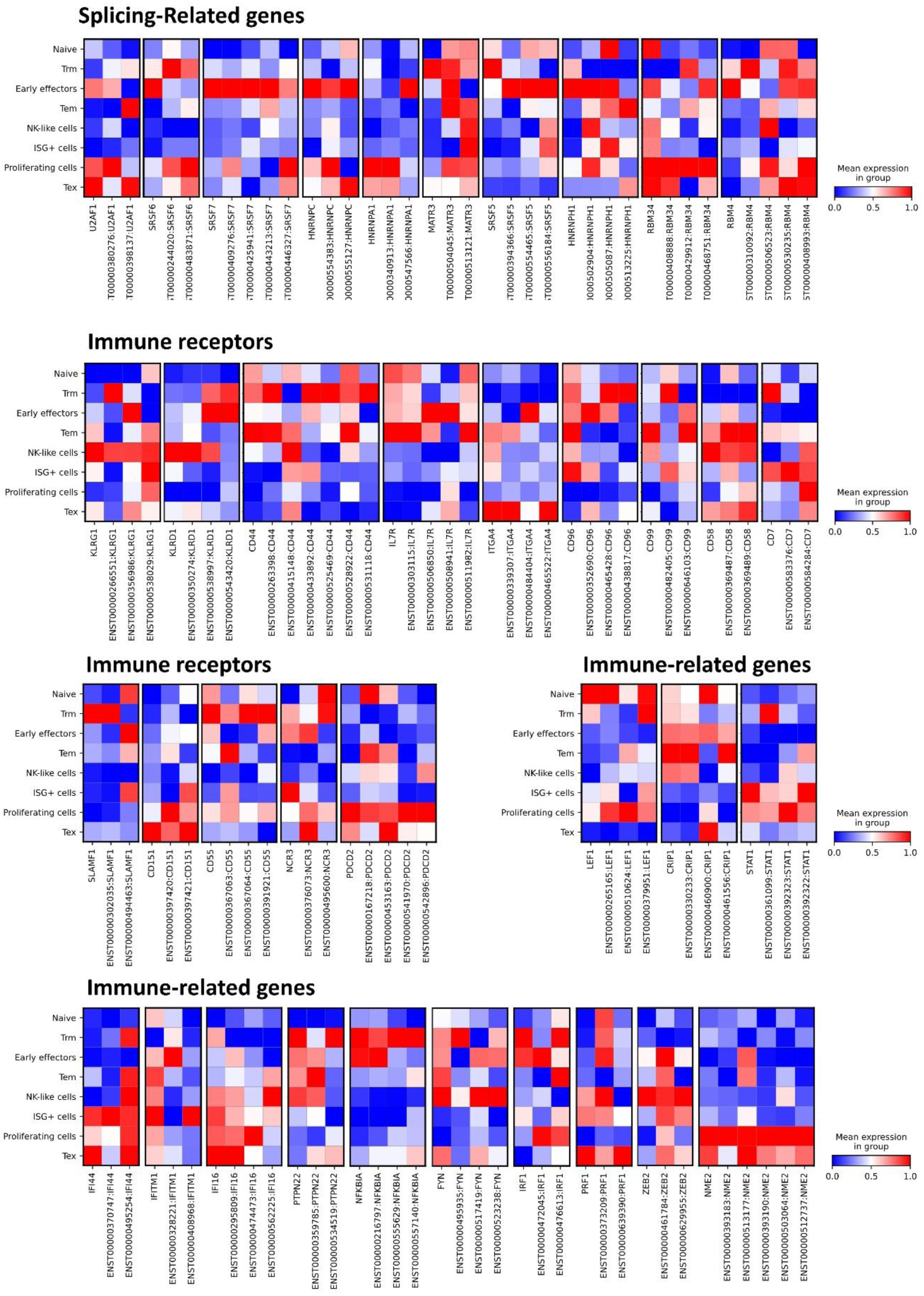
Subset-specific isoform expression across CD8⁺ T cell clusters. Heatmaps showing mean expression of selected transcript isoforms with mutually exclusive patterns across CD8⁺ T cell subsets. Expression values are min–max scaled per isoform (0 = blue; 1 = red). Isoforms were chosen from genes encoding ≥2 transcripts exhibiting cluster-restricted abundance (see Supplementary Table 3) and are arranged into three functional categories: **Splicing factors** (top panels), **Immune receptors** (middle panels), **Immune-related genes** (bottom panels).

**Extended Data Figure 4:**
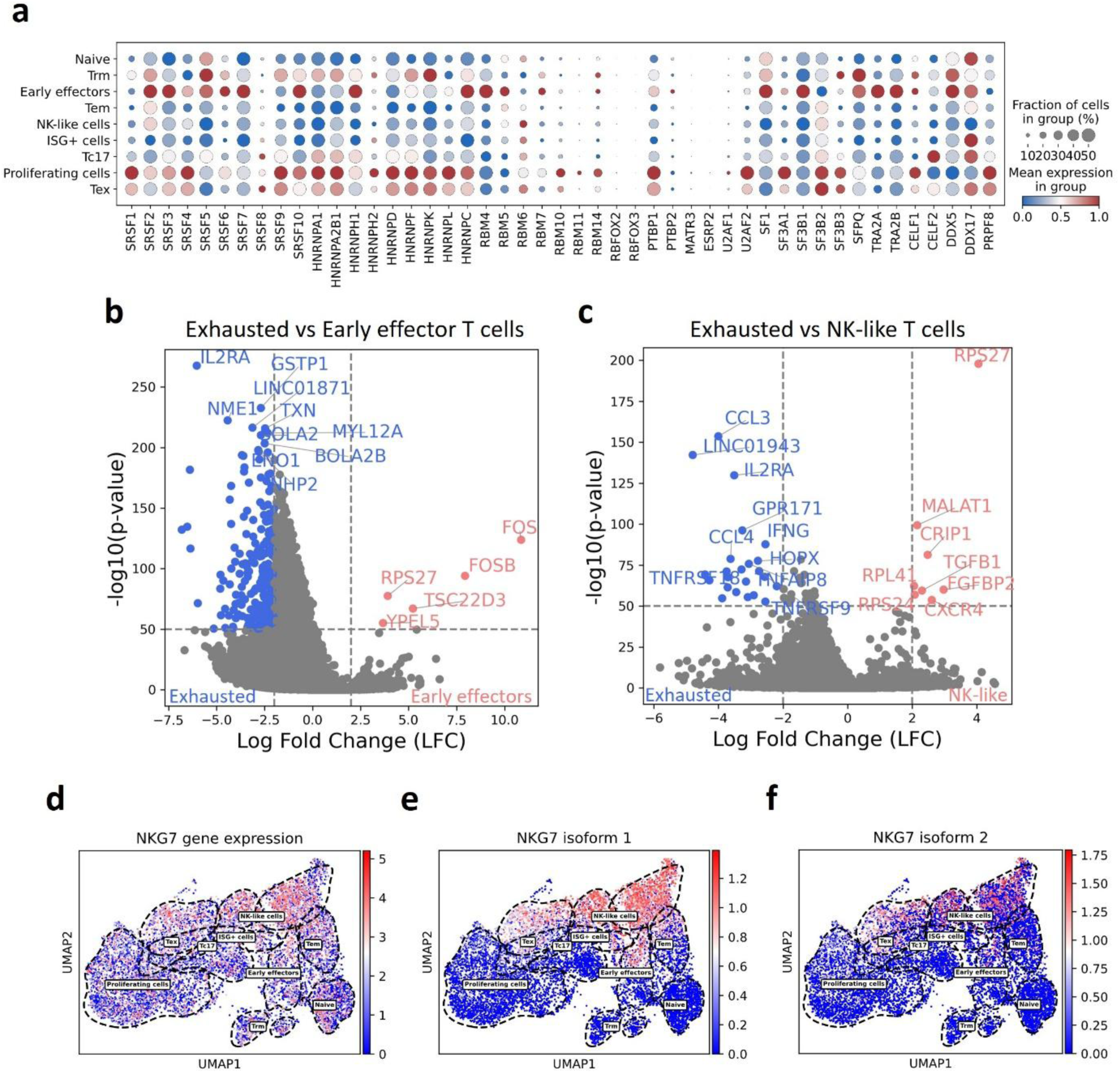
Splicing-factor expression and isoform-level regulation underpinning CD8⁺ T cell exhaustion. **a**, Dot plot of splicing-factor gene expression across CD8⁺ T cell subsets. Dot size denotes the fraction of cells in each subset; dot color reflects the mean normalized expression level (scaled 0 = low to 1 = high). **b–c,** Volcano plots of isoform differential expression between exhausted (Tex) versus early effector **(b)** or NK-like **(c)** CD8⁺ T cells. Dashed lines mark thresholds; isoforms exceeding both thresholds are blue (Tex-enriched) or red (comparison-enriched). **d–f,** UMAP projection of CD8⁺ T cells clustered by long-read splicing profiles, colored by NKG7 expression: total gene-level abundance **(d),** isoform 1 **(e),** and isoform 2 **(f).** Color scales show normalized expression intensity.

**Extended Data Figure 5:**
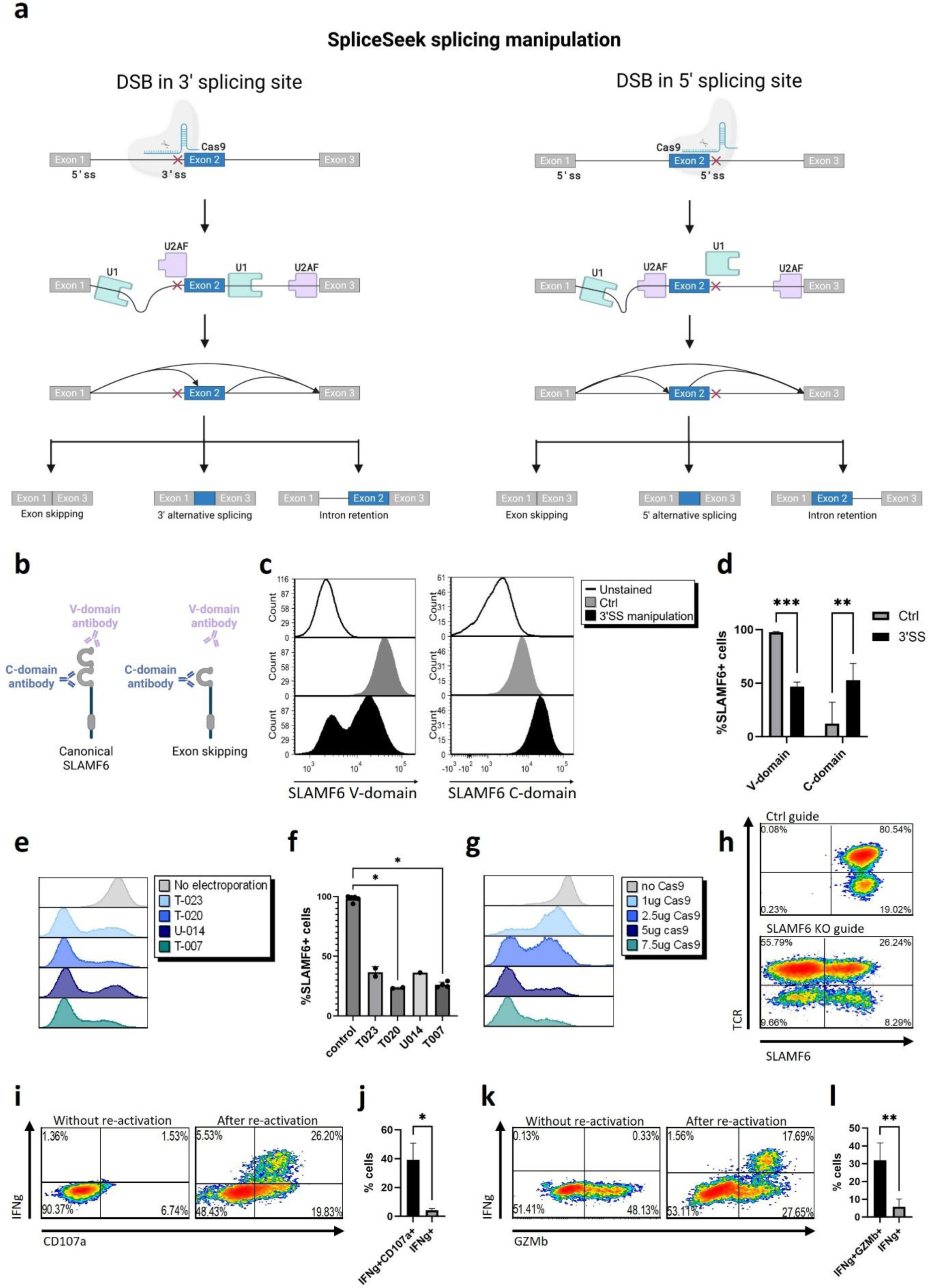
SpliceSeek CRISPR editing strategy and Cas9 electroporation optimization in primary T cells. **a,** SpliceSeek CRISPR editing strategy. Cas9 inserts indels at either the 3′ splice acceptor site (3′SS; left) or the 5′ splice donor site (5′SS; right), disrupting U1/U2AF recognition and triggering exon skipping, alternative splice-site usage or intron retention, without altering the coding sequence. Created in BioRender. **b**, Schematic of antibody epitopes: V-domain antibody (purple) binds only the canonical SLAMF6 isoform, whereas C-domain antibody (blue) recognizes all isoforms. Created in BioRender. **c**, Representative flow-cytometry histograms of Jurkat T cells edited with control or 3′SS guides and stained pan–SLAMF6 C-domain antibody (right) or a V-domain antibody specific to the canonical isoform (left). **d**, Quantification of the percentage of SLAMF6 V-domain and C-domain positive cells (mean ± SD; n = 3). **e–g**, Optimization of Cas9 electroporation parameters in primary human T cells transduced with a SLAMF6 knockout guide, assayed by SLAMF6 surface staining. **e**, Representative flow-cytometry histograms showing SLAMF6 expression after electroporation using different Lonza 2b programs. **f**, Percentage of SLAMF6⁺ cells for each electroporation program (mean ± SD, n = 3 donors). **g**, Representative histograms of SLAMF6 expression following T-007 electroporation with increasing Cas9 protein doses. **h**, Final editing efficiency under the optimized conditions (T-007 program + 5 µg Cas9): representative FACS density plots of SLAMF6 versus NY-ESO-1 TCR expression, with percentages of SLAMF6-negative cells in the SLAMF6 knockout guide condition. **i-l**, Flow cytometry of primary human T cells 6 hours after either no stimulation (left) or re-activation (right). **i, k**, Representative intracellular FACS dot plots of IFN-γ versus CD107a (i) or granzyme B (k), with quadrant percentages. **j**, Quantification of IFN-γ⁺CD107a⁺ cells (mean ± SD; n = 3). **l**, Quantification of IFN-γ⁺GZMb⁺ cells (mean ± SD; n = 4) P values were determined by two-way ANOVA with Bonferroni correction **(d),** Kruskal–Wallis test with Dunnett’s

**Extended Data Figure 6:**
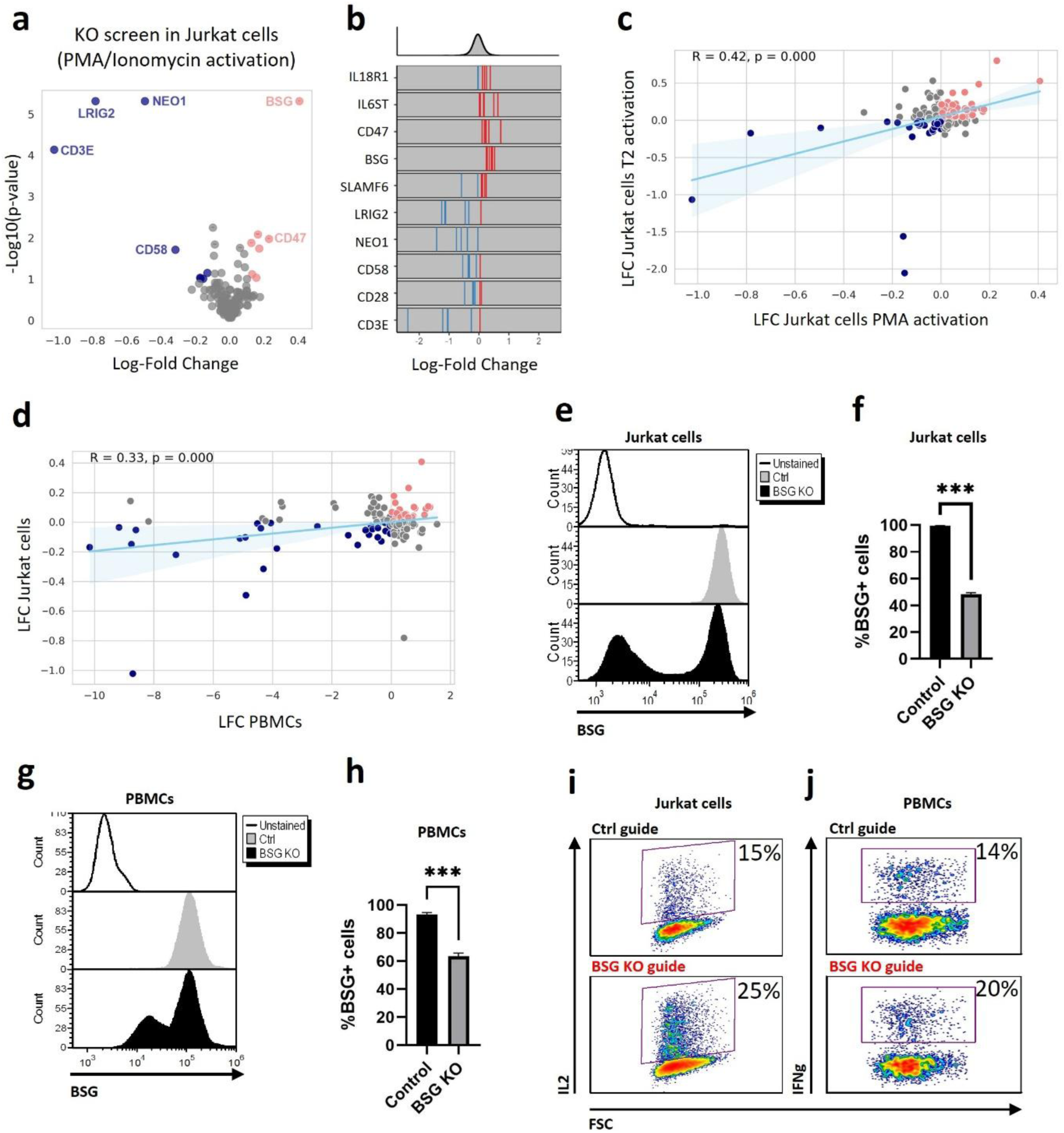
Reference CRISPR knockout screens identify BSG (CD147) as a modulator of cytokine production in T cells. **a–b**, Reference CRISPR–Cas9 knockout screen in Jurkat T cells (n = 3 independent experiments) stimulated with PMA/ionomycin and sorted into IL-2–high and IL-2–low fractions. **a,** Volcano plot of sgRNA enrichment; red - guides enriched in IL-2-high cells; blue - guides enriched in IL-2-low cells. **b,** Guide-level log₂ FC density strips for selected targets (top strip shows the overall guide distribution). **c-d**, Pearson correlation of sgRNA enrichment (log₂ FC values) in Jurkat cells activated with PMA/ionomycin versus peptide-loaded T2 cells (c) and primary PBMCs versus Jurkat PMA/ionomycin activation (d). Shaded band indicates the 95% confidence interval. **e-h** BSG (CD147) knockout efficiency validation in jurkats cells (e-f, mean ± SD; 2 different knockout guides, 3 independent experiments per guide) and PBMCs (g-h, mean ± SD; n = 3 donors). e and g are representative histograms of flow cytometry. f and h, are Quantification of BSG⁺ cells. i-j, Representative flow cytometry of cytokine expression in BSG KO cells vs controls in jurkat cells (i) and PBMCs (j). P values were determined by two-tailed paired t-test (f,h). ***P < 0.001.

**Extended Data Figure 7:**
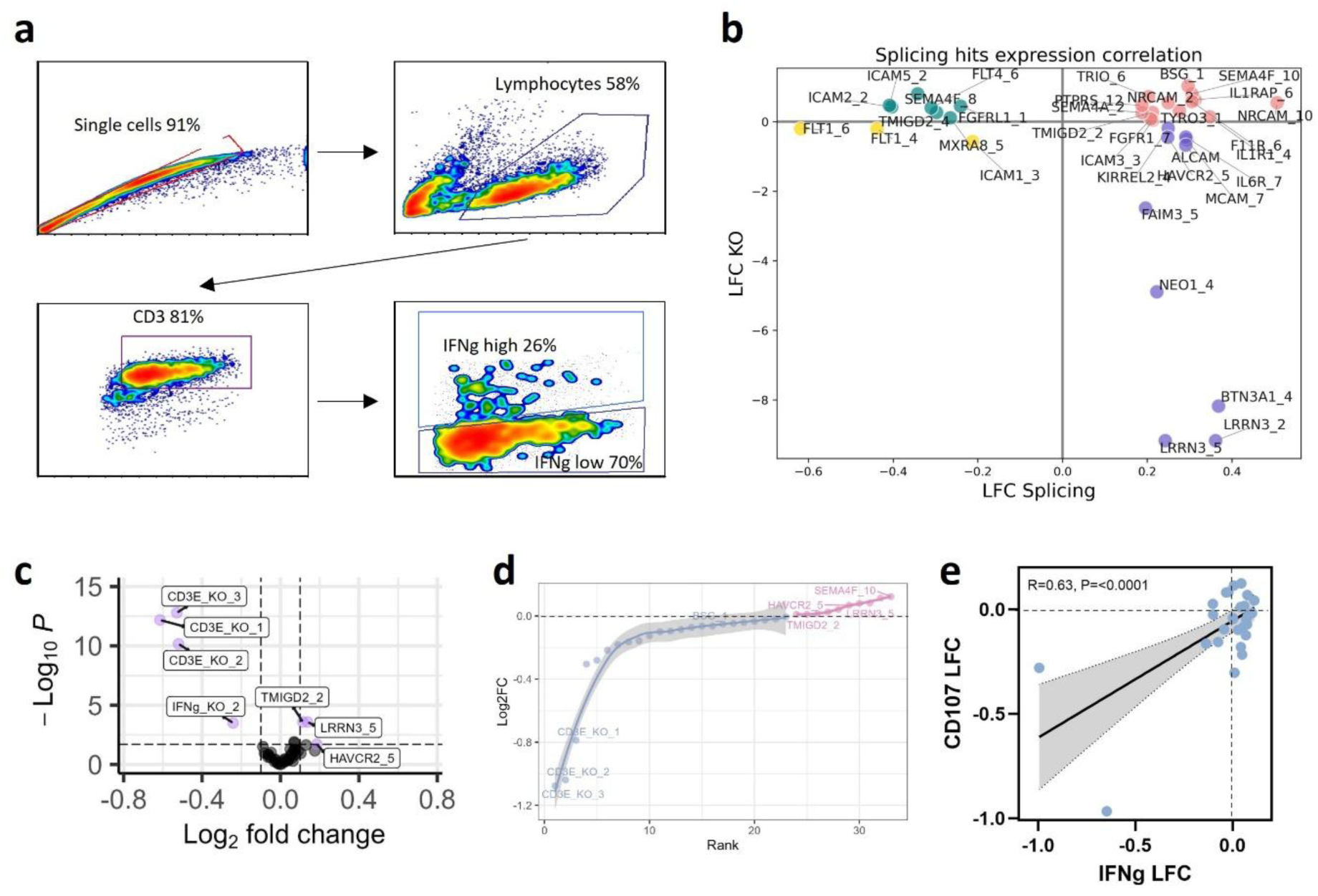
SpliceSeek sub-library screens of CD107a and IFN-γ in primary human T cells. **a,** Flow-cytometry gating strategy for the SpliceSeek screen. Singlets were first gated on FSC-A versus FSC-H to eliminate doublet cells, followed by gating for lymphocytes on FSC-A versus SSC-A. CD3⁺ T cells were then selected, and IFN-γ–high and –low populations were defined by intracellular IFN-γ staining. **b**, Comparison of LFC of isoform manipulation versus LFC of gene knockout of the top hits from the SpliceSeek screen. **c**, Volcano plot of the focused sub-library screen (24 top SpliceSeek candidates and controls; n = 9 replicates). **d**, Guide-level log₂ fold-change (CD107a-high / low) from the SpliceSeek sub-library screen in primary T cells (n = 3 donors). Guides are ordered left to right by increasing enrichment in the CD107a-high. **e**, Correlation of guide-level log₂ FC between the CD107a-high/low (x-axis) and IFN-γ-high/low (y-axis) sub-library screens. Pearson’s R and p-value are indicated, with the shaded area representing the 95% confidence interval.

**Extended Data Figure 8:**
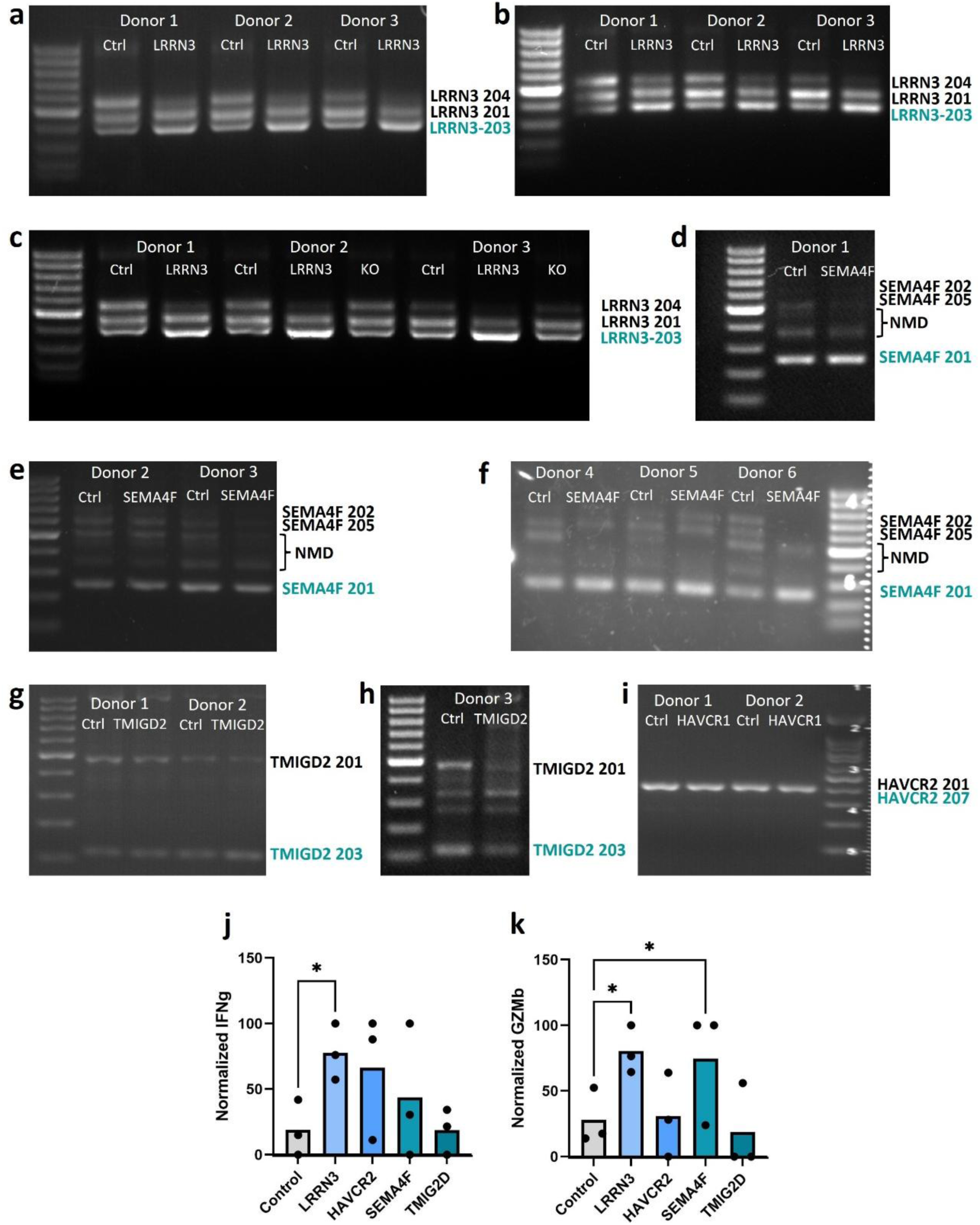
SpliceSeek hits reprogram transcript usage and enhances T cell effector functions. **a–c**, RT–PCR of LRRN3 isoforms in primary T cells from three healthy donors transduced with non-targeting control (Ctrl) or LRRN3 splice-shift guide. **d–f,** RT–PCR of SEMA4F in six donors transduced with control or SEMA4F splice-shift guides. **g–h,** RT–PCR of TMIGD2 isoforms after control or TMIGD2 splice-shift editing in three donors. **i,** RT–PCR of HAVCR2 isoform in cells edited with control or HAVCR2 splice-shift guide. **j,** IFN-γ secretion by ELISA in NY-ESO-1 TCR+ T cells (n = 3 donors, each in triplicate) transduced with control or splice-shift guides and co-cultured overnight with peptide-loaded T2 cells (0.25 µg/mL). **k,** Granzyme B secretion by ELISA in NY-ESO-1 TCR+ T cells (n = 3 donors, each in triplicate) transduced with control or splice-shift guides and co-cultured overnight with peptide-loaded T2 cells (0.5 µg/mL). Data in (j–k) was normalized by min–max scaling of donor-wise mean measurements. All values are presented as mean, with dots as donors. P values were determined by mixed-effects model with Dunnett’s multiple-comparisons test (j) and two-way ANOVA model with Dunnett’s multiple-comparisons test (k). *P < 0.05.

**Extended Data Figure 9:**
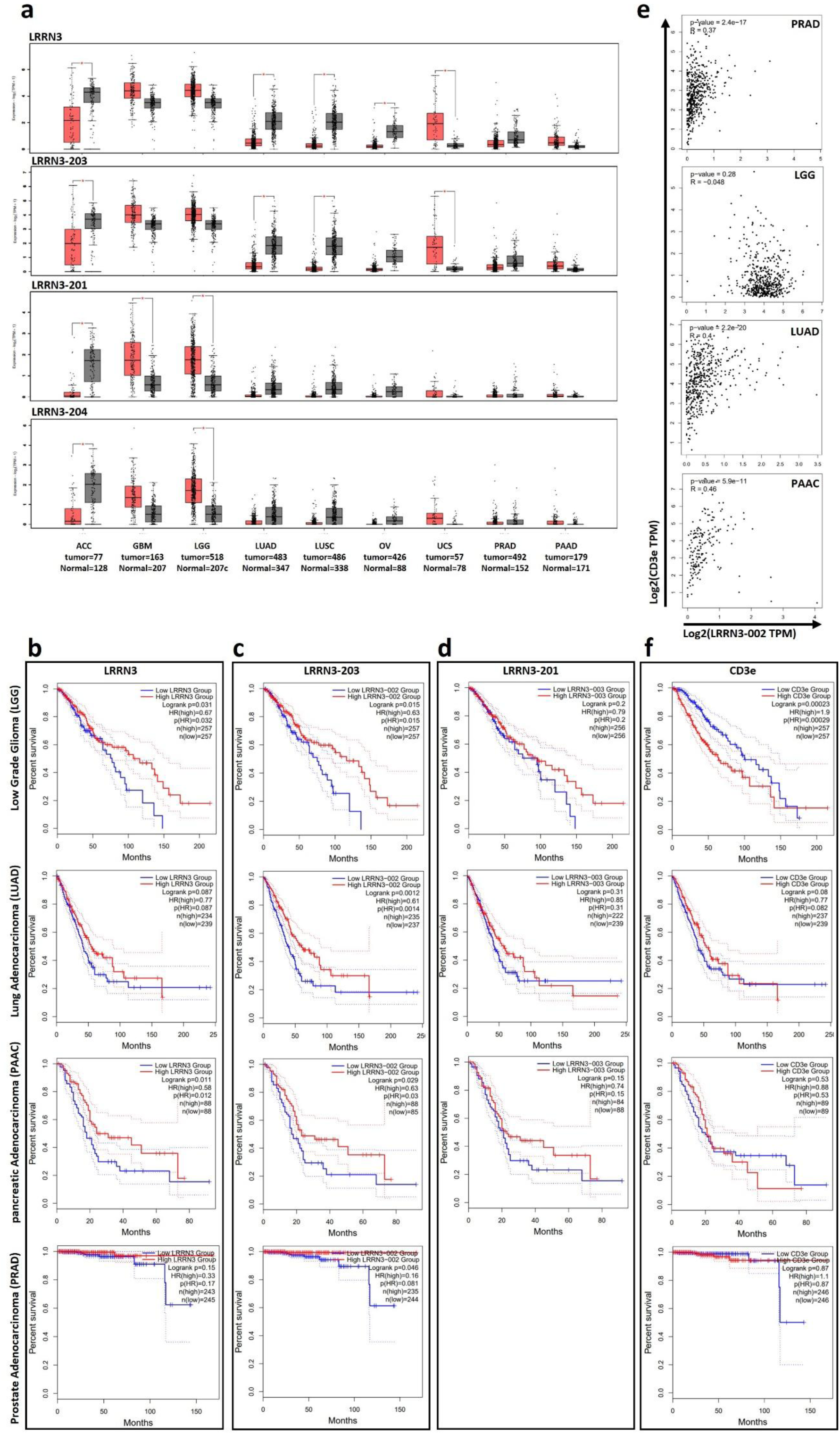
LRRN3-203 isoform is an independent favorable prognostic marker across multiple cancers. **a**, Boxplots showing expression of total LRRN3 and its three 5′-UTR isoforms (203, 201, 204) in matched healthy (gray) versus tumor (pink) tissues. **b-d**, Kaplan–Meier overall survival curves for patients with lower-grade glioma (LGG), lung adenocarcinoma (LUAD), pancreatic adenocarcinoma (PAAC) and prostate adenocarcinoma (PRAD), stratified by tumor expression of total LRRN3 (a), the LRRN3-203 isoform (b), or the LRRN3-201 isoform (c). **e**, Correlation between LRRN3- 203 and CD3E expression (log₂ TPM) across the four tumor cohorts (LGG, LUAD, PAAC, PRAD). **f**, Kaplan–Meier overall survival curves stratified by tumor expression of total total CD3E. P values were determined by log-rank test (a–d), Spearman’s rank correlation (e) and one-way ANOVA (f). *P < 0.05.

**Extended Data Figure 10:**
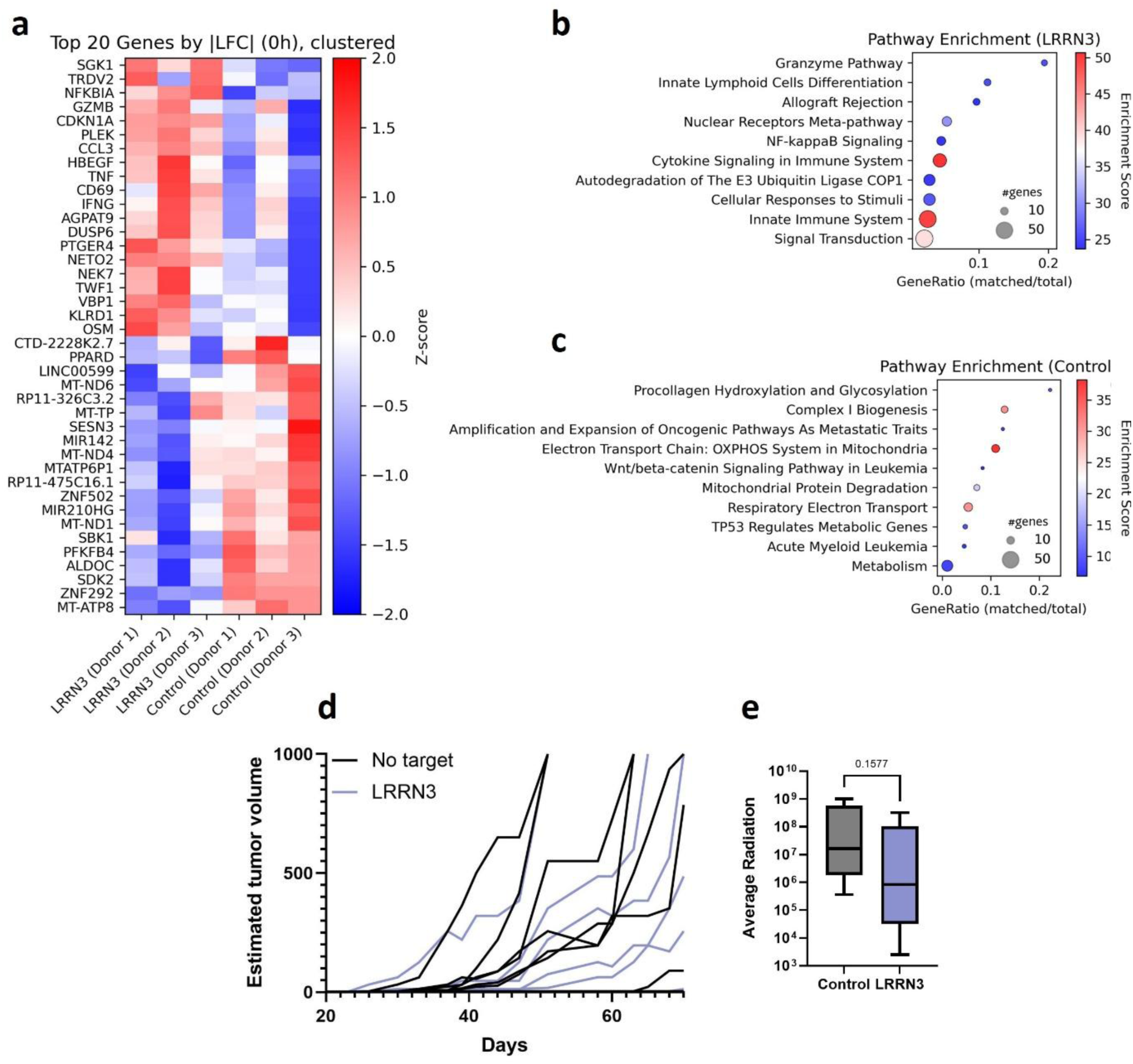
LRRN3-203 splice-shift remodels the effector transcriptome and enhances antitumor activity. **a**, Heatmap of Z-score–normalized expression for the top differentially expressed genes in LRRN3-shift versus control cells across all donors. **b-c,** Gene-set enrichment analysis of pathway enrichment in LRRN3-203 splice-shift (b) or control (c) cells. Dot size denotes the number of overlapping genes in each pathway; dot color reflects enrichment score. **d-e,** In vivo tumor control in NSG mice (n = 10 per group) co-injected subcutaneously with luciferase-expressing A375 melanoma cells and T cells transduced with either non-targeting control or LRRN3-203 splice-shift guides. **d**, Spider plot of individual tumor volume trajectories. **e**, Box-and-whisker plot of bioluminescent signal at day 60.

